# Meiotic cohesion requires Sirt1 and preserving its activity in aging oocytes reduces missegregation

**DOI:** 10.1101/2025.03.12.642822

**Authors:** Zihan Meng, Nicholas G. Norwitz, Sharon E. Bickel

## Abstract

Chromosome segregation errors in human oocytes increase dramatically as women age and premature loss of meiotic cohesion is one factor that contributes to a higher incidence of segregation errors in older oocytes. Here we show that cohesion maintenance during meiotic prophase in Drosophila oocytes requires the NAD^+^-dependent deacetylase, Sirt1. Knockdown of Sirt1 during meiotic prophase causes premature loss of arm cohesion and chromosome segregation errors. We have previously demonstrated that when Drosophila oocytes arrest and age in diplotene, segregation errors increase significantly. By quantifying acetylation of the Sirt1 substrate H4K16 on oocytes chromosomes, we find that Sirt1 deacetylase activity declines markedly during aging. However, if females are fed the Sirt1 activator SRT1720 as their oocytes age, the H4K16ac signal on oocyte DNA remains low in aged oocytes, consistent with preservation of Sirt1 activity during aging. Strikingly, age-dependent segregation errors are significantly reduced if mothers are fed SRT1720 while their oocytes age. Our data suggest that maintaining Sirt1 activity in aging oocytes may provide a viable therapeutic strategy to decrease age-dependent segregation errors.

## INTRODUCTION

Aging cells face several challenges, including (but not limited to) altered signaling pathways, changes in metabolism, mitochondrial dysfunction and oxidative damage (Lopez-Otin *et al*, 2013; Mihalas *et al*, 2024). Human oocytes initiate meiosis in the fetal ovary, arrest before birth and resume meiosis upon ovulation, which can occur decades later. Therefore, in humans, oocytes undergo years of aging before they complete the first meiotic division. As women progress through their thirties, the risk that meiotic chromosome segregation errors in aging oocytes will lead to an aneuploid pregnancy increases exponentially, a phenomenon termed the maternal age effect.

Premature loss of meiotic sister chromatid cohesion is one factor that contributes to the maternal age effect (Charalambous *et al*, 2023; Greaney *et al*, 2018; Park *et al*, 2021; Wartosch *et al*, 2021). In both meiotic and mitotic cells, cohesin-mediated linkages between sister chromatids are established during S phase and are essential for accurate chromosome segregation. During meiosis, cohesion along the arms of sister chromatids also keeps a crossover bivalent physically connected and is required for proper segregation of homologs during anaphase I. Age-dependent loss of arm or pericentric cohesion in oocytes can lead to segregation errors during the first or second meiotic division, respectively.

The mechanisms that lead to premature loss of cohesion in aging oocytes are not well-defined. One protein that helps protect aging cells by regulating cellular homeostasis and stress resistance is Sirt1/Sir2 (<u>S</u>ilent information regulator), the founding member of the highly conserved sirtuin family (Chang & Guarente, 2014; Wu *et al*, 2022). Originally identified as a chromatin silencing factor in yeast, Sirt1 is an NAD^+^-dependent deacetylase with a diverse array of substrates that regulate a wide range of physiological processes including sugar and lipid metabolism, oxidative stress, inflammation, and cellular senescence (Chen *et al*, 2020; Grabowska *et al*, 2017; McBurney *et al*, 2013; Stunkel & Campbell, 2011; Tatone *et al*, 2018; Wu *et al*., 2022). Aging is accompanied by decreased levels of Sirt1 protein and/or activity in multiple mammalian cell types, including oocytes (Di Emidio *et al*, 2014; Gong *et al*, 2014; Ma *et al*, 2015; Zhang *et al*, 2016), and this decline is considered a key factor in health challenges that become more prevalent with aging (Chen *et al*., 2020; Tatone *et al*., 2018; Wu *et al*., 2022). Because of its critical role in healthy aging, Sirt1 has become a target for therapeutic interventions in the last two decades and several small molecule Sirt1 activators have been developed and tested with promising outcomes (Dai *et al*, 2018; Grabowska *et al*., 2017; Hubbard & Sinclair, 2014; Sinclair & Guarente, 2014; Tatone *et al*., 2018).

Here we investigate the function of Sirt1 in Drosophila oocytes and provide evidence that Sirt1 activity during meiotic prophase is required for accurate chromosome segregation. Knockdown of Sirt1 during prophase I causes premature loss of arm cohesion and missegregation of recombinant homologs during meiosis I. We previously developed an experimental procedure to age Drosophila oocytes in vivo and demonstrated that when diplotene oocytes undergo aging, the incidence of meiotic segregation errors increases significantly compared to non-aged oocytes (Perkins *et al*, 2019; Subramanian & Bickel, 2008). Using this aging method, we show here that aging causes a significant decrease in Sirt1 activity in Drosophila oocytes. However, Sirt1 activity in aged oocytes is preserved if mothers are fed the Sirt1 activator, SRT1720, during the aging regimen. Furthermore, SRT1720 feeding significantly suppresses age-dependent segregation errors in Drosophila oocytes. Our results suggest that a nutritional supplementation strategy that preserves Sirt1 activity in aging oocytes might provide a viable therapeutic approach to attenuate the maternal age effect in humans.

## RESULTS

### Knockdown of Sirt1 in prophase oocytes causes a significant increase in meiotic segregation errors

To determine whether accurate chromosome segregation in Drosophila oocytes depends on Sirt1 activity during meiotic prophase, we measured segregation errors (% NDJ) in control and Sirt1 knockdown (KD) oocytes. Use of the matα-GAL4-VP16 driver (hereafter matα driver) permits us to induce expression of a short hairpin exclusively in the female germline (Januschke *et al*, 2002). In addition, because expression of this driver begins in mid-prophase (Haseeb *et al*, 2024a; Weng *et al*, 2014), this strategy allows us to investigate the effect of Sirt1 knockdown specifically on the maintenance of meiotic cohesion **(see Fig EV1A)**.

In our X*-*chromosome segregation assay, we can recover and distinguish progeny resulting from normal or aneuploid gametes and calculate the frequency of chromosome segregation errors **(Fig EV1C)**. We compared NDJ in control and Sirt1 KD oocytes that were also heterozygous for the *mtrm^KG^* allele **(Fig EV1B),** a sensitized background that results in weakened meiotic cohesion (Bonner *et al*, 2020; Haseeb *et al*, 2024b). Haploinsufficiency for *mtrm* also disables the achiasmate segregation system that operates in Drosophila oocytes in which pericentric heterochromatin-mediated association of homologs ensures proper segregation of bivalents that lack a crossover as well as crossover bivalents for which premature loss of arm cohesion causes chiasma destabilization (Dernburg *et al*, 1996; Harris *et al*, 2003; Hawley *et al*, 1992; Karpen *et al*, 1996). Therefore, via different mechanisms, heterozygosity for *mtrm^K^*^G^ renders our NDJ assay more sensitive for segregation defects that arise due to premature loss of cohesion.

To induce knockdown, we utilized a previously described recombinant *mtrm^KG^* matα driver chromosome that results in robust Gal4 expression (Haseeb *et al*., 2024a; Perkins *et al*, 2016). Segregation errors were significantly greater in KD oocytes (*mtrm*, matα driver ➔ short hairpin) than control oocytes (*mtrm*, no driver ➔ short hairpin). Two different Sirt1 hairpins yielded similar results **(Fig 1A)**. These results demonstrate that Sirt1 function during meiotic prophase is essential for accurate chromosome segregation in Drosophila oocytes.

**Figure 1.**
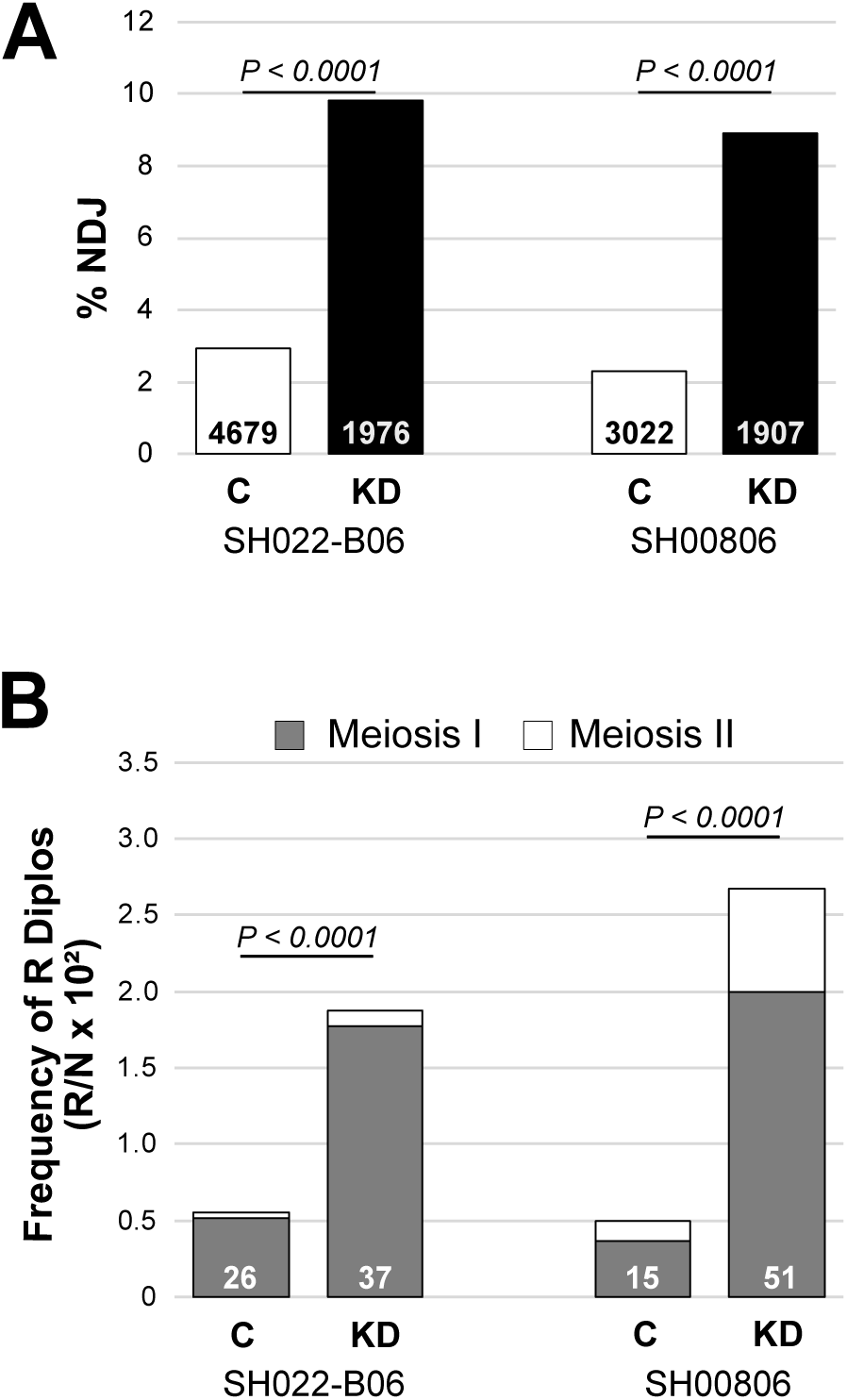
Sirt1 knockdown (KD) in oocytes during prophase causes chromosome segregation errors. **A.** Meiotic segregation errors (% NDJ) are presented for two different Sirt1 short hairpins (SH022-B06 & SH00806). Knockdown of Sirt1 during meiotic prophase (**black**, *mtrm,* matα driver **→** hairpin) significantly increases NDJ compared to each respective Control (**white**, *mtrm,* no driver **→** hairpin). See also Figure EV1. The number of flies scored (N) is provided in the bar for each genotype. **B.** Diplo-X females from the NDJ tests in Fig 1A were used in a subsequent test to determine the recombinational history of missegregating chromosomes (see Methods and Figure EV1). For both Sirt1 hairpins, KD significantly increases the missegregation of recombinant homologs, consistent with premature loss of arm cohesion. The number of recovered Diplo-X females with at least one recombinant X chromosome (R Diplo) is indicated at the bottom of each bar. The graph presents the number of R Diplos divided by the total number of progeny in the NDJ test (N) and multiplied by 100. Gray represents Diplo-X females that inherited two homologs (Meiosis I error) and white depicts those that inherited two sisters (Meiosis II error). P values shown compare Meiosis I errors in Sirt1 KD and control oocytes.

Using Diplo-X progeny from the NDJ tests described above, we performed an additional cross that allowed us to determine if Sirt1 KD caused recombinant homologs to missegregate at a higher frequency **(Fig EV1D)**. Because sister chromatid cohesion distal to a crossover keeps recombinant homologs associated (Bickel *et al*, 2002; Buonomo *et al*, 2000; Hodges *et al*, 2005), premature loss of arm cohesion will allow a recombinant bivalent to segregate randomly during the first meiotic division. Therefore, increased missegregation of recombinant homologs in Sirt1 KD oocytes would support the hypothesis that maintenance of arm cohesion during meiotic prophase relies on Sirt1.

Because the females we used in the NDJ test were heterozygous for recessive visible markers **(Fig EV1B),** we could deduce the X chromosome genotype of each Diplo-X female (arising from missegregation) by phenotyping her sons **(Fig EV1D).** Specifically, we could determine 1) whether the missegregating X chromosomes had undergone recombination before missegregation and 2) based on the centromere-proximal marker *car*, we could determine if the two chromosomes are homologs (*car^+/-^)* or sisters (*car^+/+^* or *car^-/-^)*, indicating missegregation at Meiosis I or Meiosis II, respectively. For each of the Sirt1 short hairpins, knockdown significantly increased the frequency at which recombinant homologs missegregated (Meiosis I), consistent with premature loss of arm cohesion allowing their separation prior to anaphase I **(Fig 1B)**. For one hairpin (SH00806), we also observed a significant increase in the frequency at which Diplo-X females inherited two sister chromatids from a crossover bivalent (P=0.003) consistent with premature loss of pericentromeric cohesion. However, because some of our SH00806 stocks appeared unstable, we continued our experiments using solely the SH022-B06 hairpin.

### Cohesion maintenance in prophase oocytes depends on Sirt1

To assay directly whether cohesion is lost prematurely when Sirt1 is knocked down during meiotic prophase, we performed FISH (Fluorescence In Situ Hybridization) on mature *Sirt1^SH022-B06^* KD and control oocytes that were wild-type for *mtrm*. We quantified cohesion defects using two differently labeled X-chromosome probes **(Fig 2A)**; one hybridizes to a large block of heterochromatin near the centromere and the other recognizes a 100Kb region on the distal arm. **Fig 2B** provides examples of oocytes in which arm cohesion is intact (one or two arm spots, top images) and those that we score as cohesion defective (three or four arm spots, bottom images). In matα ➔ Sirt1 KD oocytes, arm cohesion defects were significantly higher than in control oocytes. **Fig 2C** presents data from two independent replicates. Our scoring did not uncover any pericentric cohesion defects in Sirt1 KD or control oocytes. However, the large size of the satellite repeat (11Mb) recognized by this probe may hamper detection of cohesion defects near the centromere of the X chromosome (Haseeb *et al*., 2024a; Haseeb *et al*., 2024b). The finding that arm cohesion is disrupted when Sirt1 is knocked down indicates that cohesion maintenance in Drosophila oocytes during meiotic prophase depends on Sirt1 protein.

**Figure 2.**
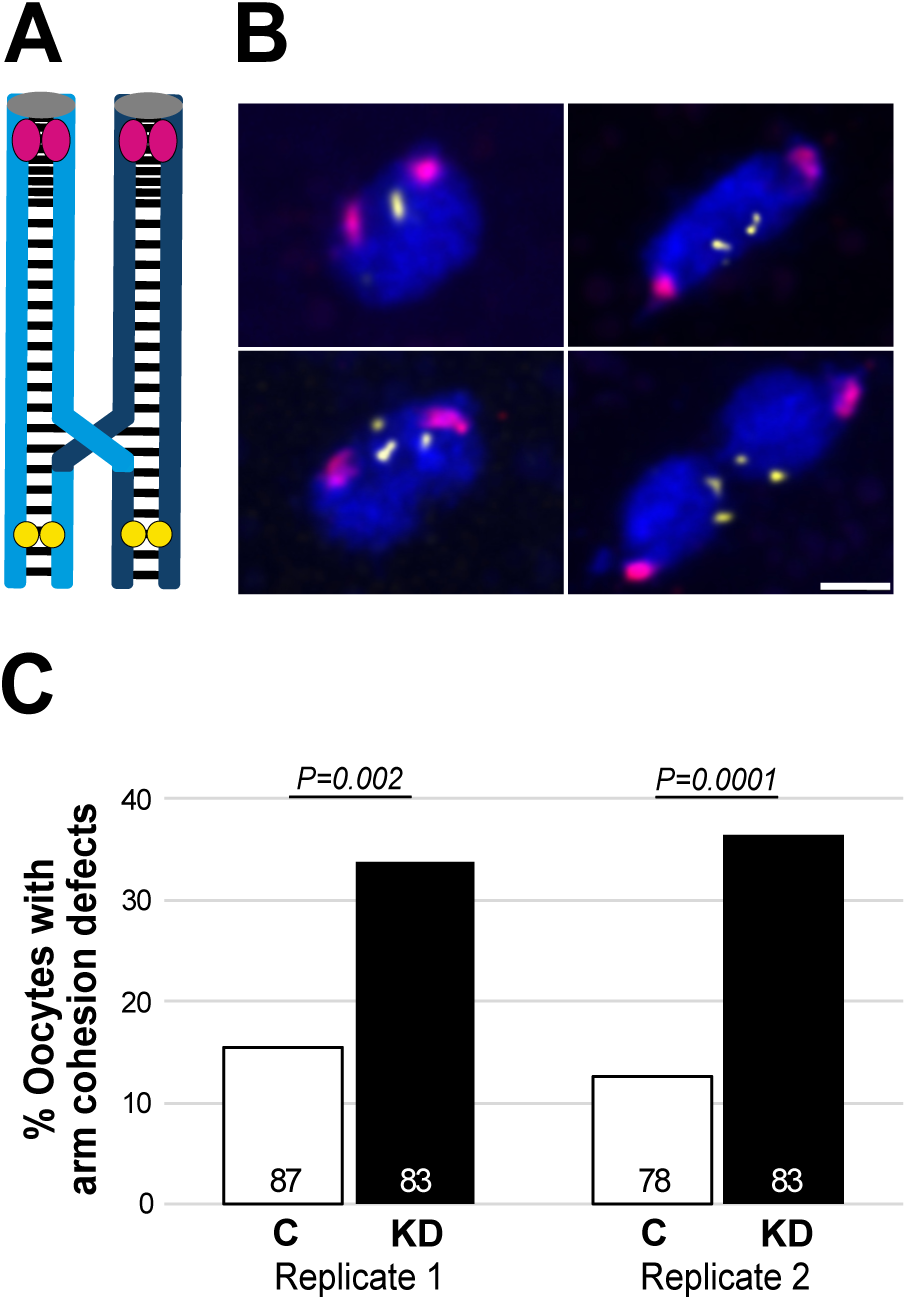
Arm cohesion defects increase significantly in Sirt1 KD oocytes. **A.** Drosophila X chromosome bivalent with a single crossover is shown. Light blue and dark blue sister chromatids are held together by cohesion, depicted as black lines. OligoPaint probe (yellow) hybridizes to a distal location on the X chromosome arm and the 359 bp satellite repeat probe (magenta) hybridizes close to the centromere (gray). **B.** Top images show oocyte chromosomes with intact arm cohesion (one or two yellow spots). Examples of premature loss of arm cohesion (three or four yellow spots) are provided on the bottom. Images are maximum projections of deconvolved confocal Z series. Scale bar, 2 µm. **C.** The percentage of Sirt1 control (white) and KD (black) oocytes with arm cohesion defects are shown for two independent replicates. Knockdown of Sirt1 during meiotic prophase significantly increases the percentage of oocytes with arm cohesion defects. The number of oocytes scored is indicated within each bar. No pericentric cohesion defects were detected in either replicate. Note that both Control and KD genotypes are wild-type for *mtrm*.

### Aging reduces Sirt1 activity on oocyte chromosomes

Given that reduction of Sirt1 protein and/or activity has been shown to accompany aging in multiple cell types (Di Emidio *et al*., 2014; Gong *et al*., 2014; Ma *et al*., 2015; Zhang *et al*., 2016) we next investigated whether Drosophila oocytes suffer a decline in Sirt1 protein and/or activity when they undergo aging. We have previously described an experimental strategy to age Drosophila oocytes in vivo using a sensitized genetic background in which functional cohesin is reduced by half (*smc111* /+) and the achiasmate segregation system is disabled (*mtrm^KG^/+*). In this genotype, cohesion is weakened but not eliminated (Subramanian & Bickel, 2008). When *mtrm^KG^/smc1Δ* females were subjected to our aging regimen **(Fig EV2)**, segregation errors were significantly higher in aged oocytes than in non-aged oocytes (Perkins *et al*., 2019; Subramanian & Bickel, 2008). Moreover, oocytes at stages 7 and 8, which arrest and age in diplotene, are the most vulnerable to age-dependent segregation errors (Subramanian & Bickel, 2008) and this is the meiotic stage at which human oocytes arrest for decades. For the studies described below (cytology and NDJ), we utilized females that were heterozygous for a *mtrm^KG^ smc111* recombinant chromosome **(Fig EV2)**, subjected them to the aging regimen and focused on stages 7 and 8 for cytological analysis.

We first validated immunoreagents using ovaries from Sirt1 control and *sirt1* null transheterozygotes (*sirt1^4.5^/sirt1^5.26^*). Using an anti-Sirt1 monoclonal antibody to localize Sirt1 protein in Drosophila ovarioles, we observed predominantly nuclear signal in nurse cells and follicle cells as well as the oocyte nucleus **(Fig EV3).** This pronounced nuclear Sirt1 signal was absent in egg chambers from *sirt1* null females.

When we quantified the Sirt1 signal associated with oocyte DNA in control and null genotypes, we confirmed that Sirt1 signal on oocyte chromosomes was significantly reduced in null oocytes compared to control **(Fig 3A-B)**. Moreover, when the matα driver was used to induce knockdown of Sirt1 during meiotic prophase, Sirt1 signal associated with oocyte chromosomes was comparable to that observed in *sirt1* null oocytes **(Fig 3A-B)**.

**Figure 3.**
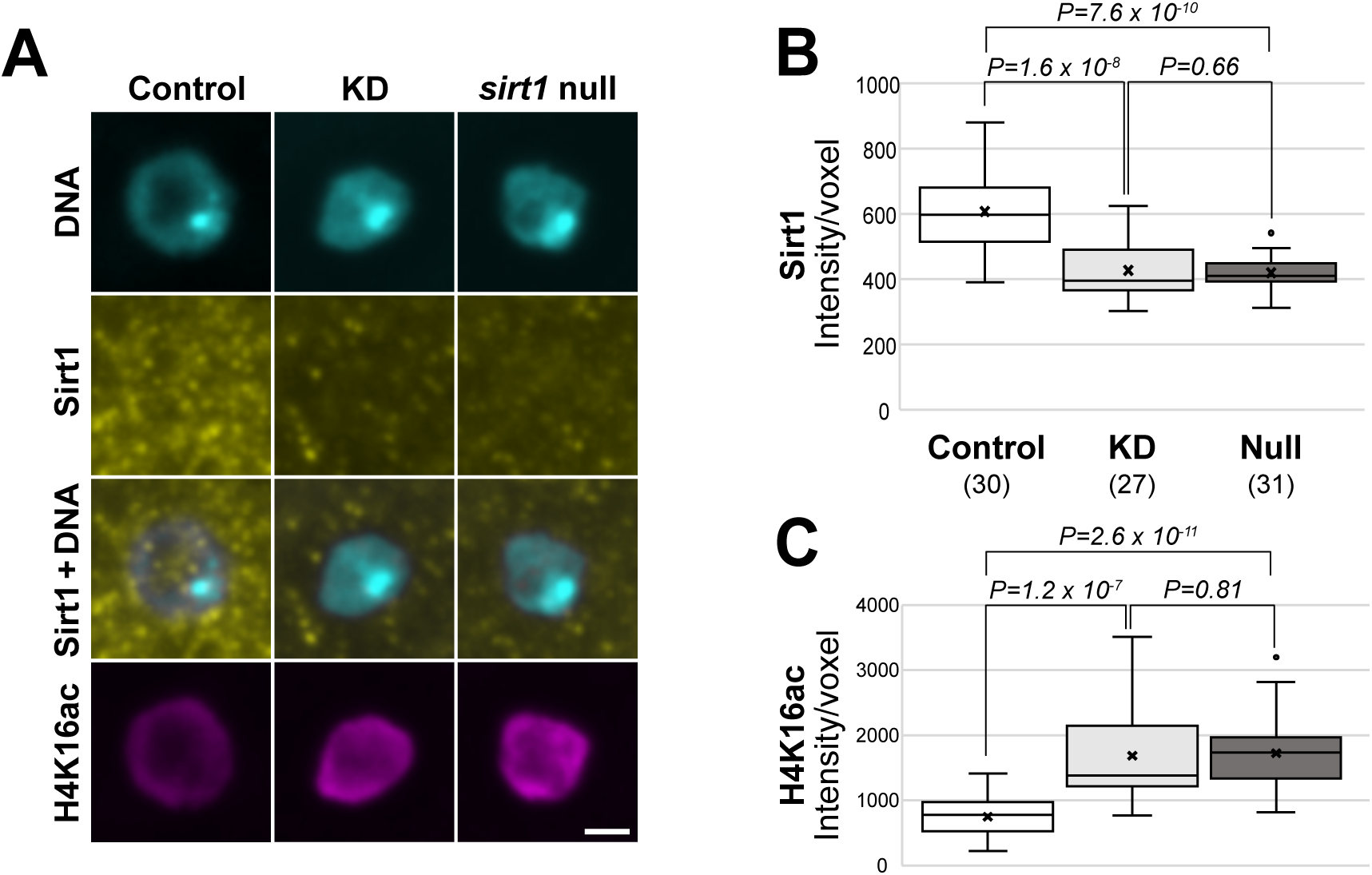
Deacetylation of H4K16 on oocyte DNA provides a readout for Sirt1 activity in vivo. **A.** Oocyte DNA (cyan) with Sirt1 (yellow) and H4K16ac (magenta) immunolocalization in Sirt1 Control, KD and *sirt1* null oocytes (*sirt1^4.5^/sirt1^5.26^*). Stage 7 oocyte chromosomes are shown as a maximum intensity projection of a confocal Z series. Scale bar, 2 μm. **B-C.** Quantification of chromosome-associated Sirt1 and H4K16ac signal intensity in stage 7-8 oocytes. The Sirt1 and H4K16ac signal intensities are inversely proportional. Compared to Control, Sirt1 signal associated with oocyte DNA is significantly lower in Sirt1 KD and *sirt1* null oocytes and H4K16ac is significantly higher in these genotypes, consistent with loss of the Sirt1 deacetylase. The number of oocytes analyzed for each genotype is shown in parentheses.

To measure Sirt1 activity in vivo, we utilized an antibody specific for histone H4 when acetylated at Lysine 16 (H4K16ac), a known Sirt1 deacetylation target (Vaquero *et al*, 2004). As previously reported by others (Samata *et al*, 2020), we observed H4K16ac signal primarily on the oocyte chromosomes of Drosophila egg chambers **(Fig EV3)**. Absence of Sirt1 enzymatic activity in *sirt1* null females significantly increased the H4K16ac signal associated with oocyte DNA compared to control oocytes **(Fig 3A,C)**. The H4K16ac signal on oocyte chromosomes was comparable in *sirt1* null and Sirt1 KD, consistent with neglible levels of Sirt1 associated with oocyte DNA in these two genotypes **(Fig A-C)**. These data confirm that the H4K16ac signal on oocyte DNA provides a reliable method to monitor Sirt1 activity in the Drosophila oocyte.

We next asked whether the level of Sirt1 protein or activity associated with oocyte chromosomes is impacted when Drosophila oocytes undergo aging. We utilized our normal aging regimen **(Fig EV2)** to generate *mtrm^KG^ smc111* / + aged and non-aged ovarioles which were fixed and processed for Sirt1 or H4K16ac immunostaining. As a control, we also fixed and stained ovaries from *sirt1* null females that were not subjected to the aging regimen.

As shown in **Fig 4A-B**, aging did not affect the amount of Sirt1 protein on oocyte DNA. However, a significant reduction of Sirt1 signal associated with oocyte chromosomes was still detected for *sirt1* null oocytes.

**Figure 4.**
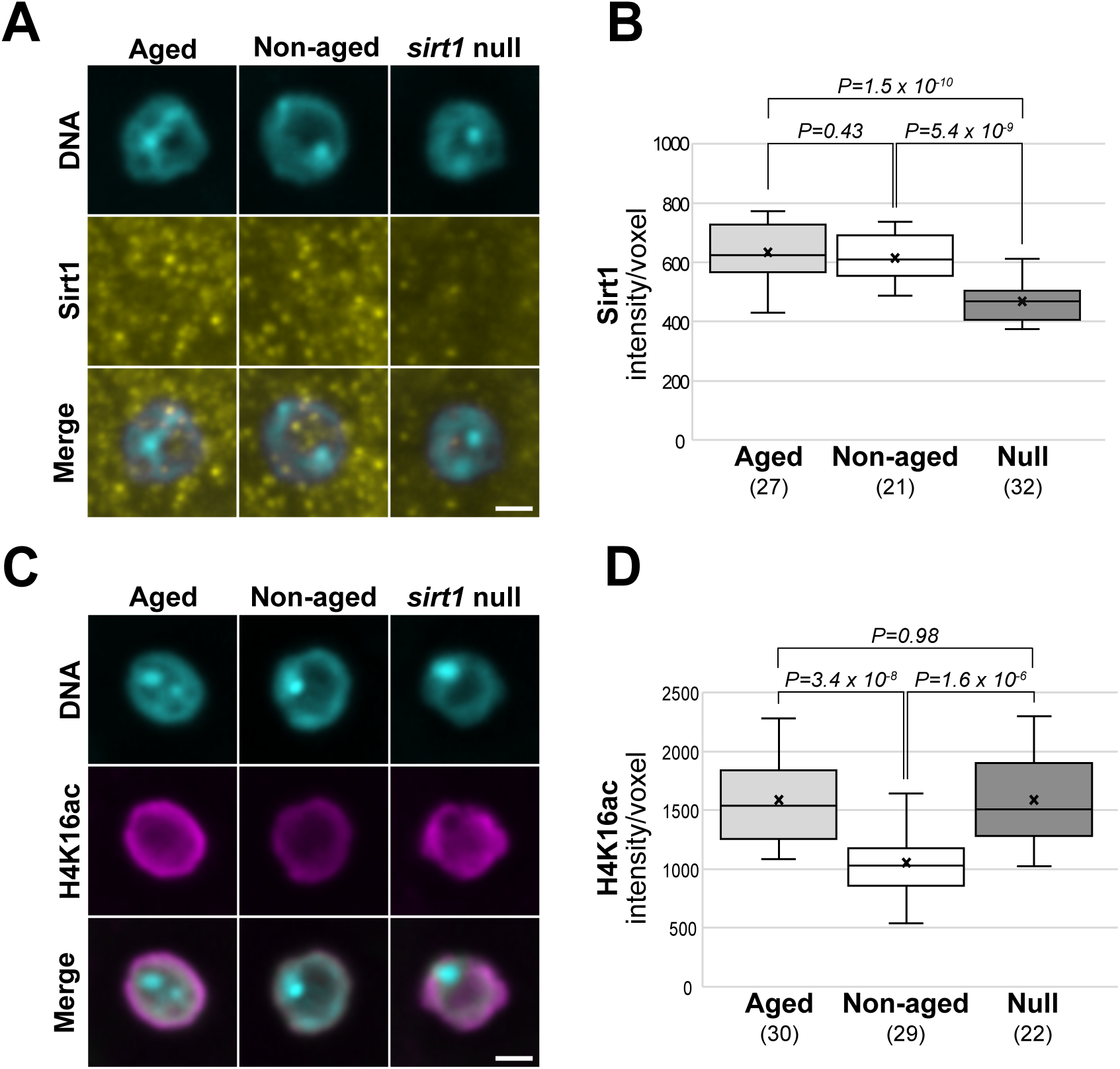
Sirt1 activity on oocyte chromosomes declines during aging. **A & C.** Sirt1 (yellow) and H4K16ac (magenta) immunostaining on the DNA (cyan) of stage 7 Aged, Non-aged and *sirt1* null oocytes. All images are maximum intensity projections of confocal Z series. Scale bar, 2 μm. **B.** Signal intensity of Sirt1 on the DNA of stage 7 & 8 oocytes indicates that chromosome-associated Sirt1 does not decline during aging. **D.** Quantification of H4K16ac signal on oocyte DNA demonstrates that aging causes a significant increase in acetylation, consistent with loss of Sirt1 deacetylase activity. The number of oocytes scored in B and D is shown in parentheses.

Although we did not detect a decrease in the amount of Sirt1 protein associated with the DNA of oocytes that had undergone aging, the H4K16ac signal was significantly higher on the oocyte DNA of aged oocytes compared to non-aged oocytes **(Fig 4C-D)**. Because a higher H4K16ac signal corresponds to decreased Sirt1 deacetylase activity, these data indicate that aging causes a decline in Sirt1 activity associated with oocyte chromosomes. Notably, acetylation of H4K16 on the chromosomes of aged oocytes was comparable to that for *sirt1* null oocytes **(Fig 4C-D)**.

### Feeding females SRT1720 prevents the decline of SIRT1 activity in aging oocytes

Several small molecule activators of Sirt1 have been developed and recent work suggests that nutritional supplementation may provide a valuable therapeutic approach for age-related pathologies (Dai *et al*., 2018; Grabowska *et al*., 2017; Hubbard & Sinclair, 2014; Sinclair & Guarente, 2014; Tatone *et al*., 2018). One such activator, SRT1720, was found to be 1000 times more effective than the naturally occurring Sirt1 activator, resveratrol (Milne *et al*, 2007). In vitro, 10µM SRT1720 was able to increase Sirt1 activity over 7-fold (Milne *et al*., 2007) and in vivo studies have demonstrated that feeding mice SRT1720 can delay and/or improve age-dependent health issues such as insulin resistance and inflammation (Milne *et al*., 2007; Minor *et al*, 2011; Mitchell *et al*, 2014). Therefore, we asked whether feeding SRT1720 to Drosophila females during the oocyte aging regimen could counteract the aging-induced decline of Sirt1 activity on oocyte chromosomes.

We carried out our standard aging regimen with two treatment groups: DMSO only and 10µM SRT1720 dissolved in DMSO **(Fig 5A)**. Fresh yeast paste was prepared daily with the addition of DMSO or SRT1720 and applied to a fresh glucose plate. At the end of the four-day aging regimen, ovaries were fixed and immunostained to quantify acetylation of H4K16 on the chromosomes of aged and non-aged oocytes at stages 7 and 8.

**Figure 5.**
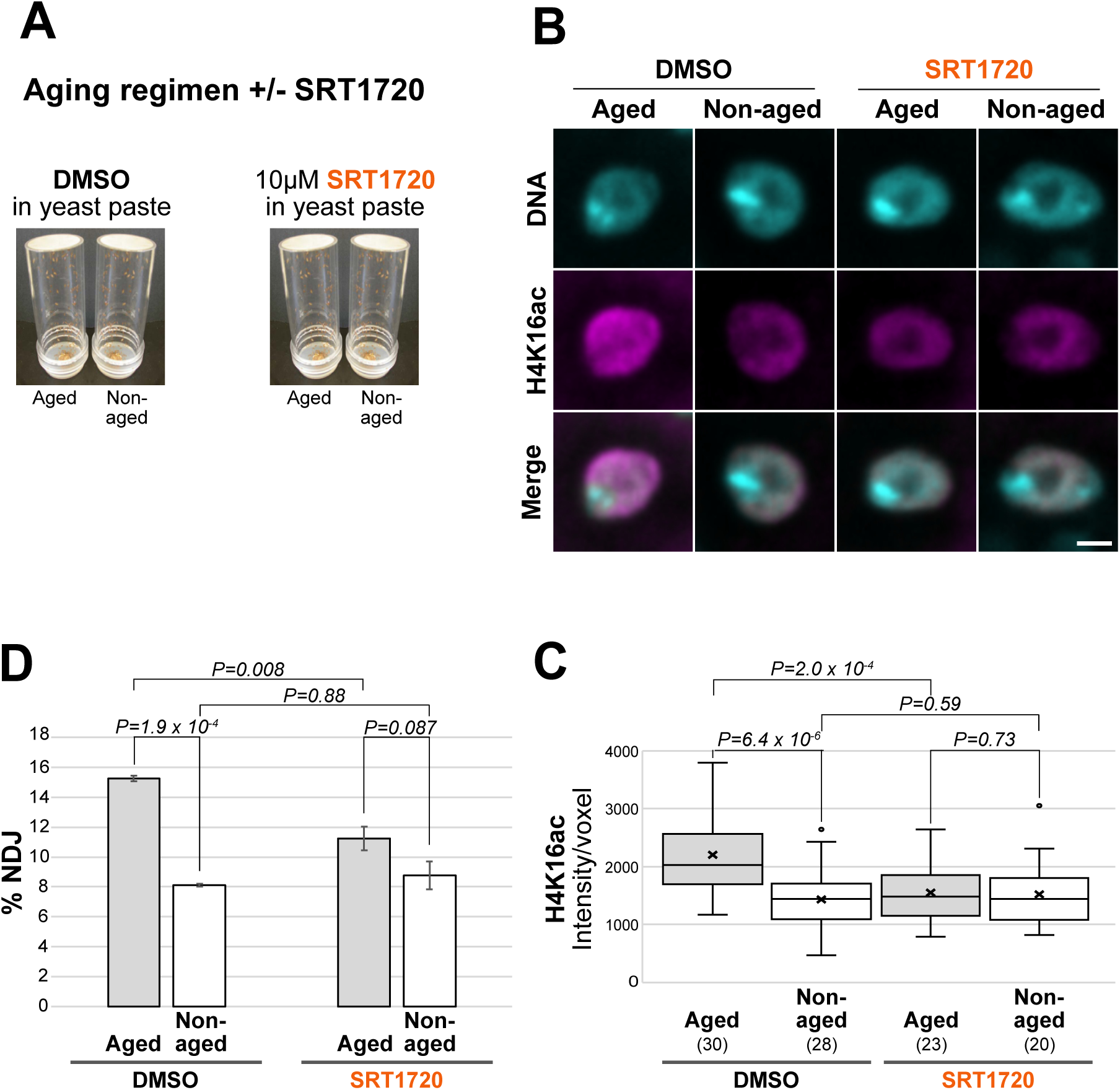
SRT1720 feeding preserves Sirt1 activity on oocyte DNA during aging and suppresses age-dependent segregation errors. **A.** Aging-regimen +/- SRT1720 feeding. **B.** H4K16ac signal (magenta) on the chromosomes of aged and non-aged oocytes (stage 7) after mothers were fed DMSO or 10µM SRT1720 during the aging regimen. All images are maximum intensity projections of confocal Z series. Scale bar, 2 μm. **C.** Quantification of H4K16ac signal intensity on oocyte DNA (stages 7 and 8) indicates that feeding mothers SRT1720 during the aging regimen prevents the significant increase in H4K16 acetylation that occurs when oocytes undergo aging in the presence of DMSO. Acetylation of H4K16 is significantly lower in aged oocytes that were exposed to SRT1720 than aged oocytes that were not (DMSO). With SRT1720 supplementation, H4K16ac signal is comparable in aged and non-aged oocytes, consistent with preservation of Sirt1 activity during aging. The number of oocytes for which H4K16ac signal was quantified is indicated in parentheses for each conditon. **D.** Chromosome segregation errors (% NDJ) were measured for aged and non-aged oocytes whose mothers were fed DMSO or SRT1720 during the aging regimen. For each condition, the mean % NDJ for three independent experiments is plotted with error bars (SEM). Compared to aged oocytes in the DMSO treatment group, SRT1720 feeding significantly reduced NDJ in aged oocytes. A significant difference between the NDJ of aged versus non-aged oocytes is not observed when mothers are fed SRT1720 during the aging regimen.

As shown in **Fig 5B-C**, aging in the absence of SRT1720 (DMSO only) caused a significant increase in the H4K16ac signal on the oocyte DNA, indicative of reduced Sirt1 activity. However, when females were fed 10µM SRT1720 during the aging regimen, acetylation of H4K16 on the chromosomes of aged oocytes was comparable to the low level observed in non-aged oocytes. These data indicate that feeding females SRT1720 can prevent the decline in Sirt1 activity that occurs when their oocytes undergo aging in the absence of this Sirt1 activator.

### Age-dependent segregation errors are suppressed when mothers are fed SRT1720 during oocyte aging

Because SRT1720 feeding prevented a decline of Sirt1 activity on oocyte chromosomes during aging, we next asked whether this supplementation could also reduce age-dependent segregation errors. We carried out the four-day oocyte aging regimen in the presence and absence of the Sirt1 activator, SRT1720, and divided the females into vials for the NDJ assay. **Fig 5D** and **Table EV2** present the data from three independent experiments. In the treatment group that lacked SRT1720 (DMSO only), meiotic segregation errors were significantly higher in aged oocytes than in non-aged oocytes, as expected. When females were fed 10µM SRT1720 during the aging regimen, NDJ in aged oocytes was slightly higher than that observed for non-aged oocytes, but the difference between these two groups was not significant. Notably, segregation errors were significantly lower for aged oocytes in the SRT1720 treatment group than for aged oocytes in the DMSO treatment group. In addition, non-aged oocytes from both treatment groups (+ or – SRT1720) exhibited similar levels of NDJ. These data demonstrate that if the aging-induced decline of Sirt1 activity on oocyte chromosomes is prevented with SRT1720 nutritional supplementation, age-dependent segregation errors can also be prevented.

## DISCUSSION

Here we demonstrate that Sirt1 is required during prophase I for accurate chromosome segregation in Drosophila oocytes. Sirt1 KD during meiotic prophase significantly increases the number of oocytes with arm cohesion defects, as evidenced by FISH (*mtrm^+^* oocytes). Premature loss of arm cohesion leading to chiasma destabilization is consistent with the increased frequency at which recombinant homologs missegregate in Sirt1 KD oocytes.

A link between Sirt1 and cohesion has previously been described in budding yeast. Cohesion within silent chromatin domains depends on the yeast ortholog, Sir2, but does not require its deacetylase activity or its silencing partners (Wu *et al*, 2011). While this mechanism may operate at a limited number of other locations within the yeast genome, it is not universally required for arm cohesion in budding yeast (Chen *et al*, 2016).

Sirt1 has a diverse set of substrates and could impact cohesion maintenance by a variety of mechanisms, either direct or indirect. Interestingly, acetylation of the cohesin loader NIPBL and the cohesion establishment factor ESCO2 are significantly elevated in Sirt1 knockout cells (Chen *et al*, 2012). Although these quantitative proteomics data cannot distinguish whether these proteins are deacetylated directly by Sirt1 or another deacetylase regulated by Sirt1, they suggest an intriguing mechanistic link between Sirt1 and cohesion. We have recently reported that newly synthesized cohesin loads onto oocyte chromosomes and generates new cohesive linkages during meiotic prophase in Drosophila oocytes, a process we have termed “cohesion rejuvenation” (Haseeb *et al*., 2024b; Weng *et al*., 2014). Moreover, the Drosophila orthologs of the cohesin loader (Nipped-B) and the establishment factor (Eco) are both required for cohesion rejuvenation in Drosophila oocytes (Haseeb *et al*., 2024b; Weng *et al*., 2014). Therefore, cohesion defects in Sirt1 KD oocytes may arise because the rejuvenation process is compromised by loss of Sirt1 activity.

Sirt1 also modulates the activity of several transcription factors (Chen *et al*., 2020; Grabowska *et al*., 2017; McBurney *et al*., 2013; Stunkel & Campbell, 2011; Wu *et al*., 2022) that could impact meiotic cohesion via an indirect manner such as controlling oxidative stress within the oocyte. Sirt1 positively regulates the expression or activity of several antioxidant enzymes (Alam *et al*, 2021; Li, 2014; Singh *et al*, 2018) and we have previously shown that induction of oxidative stress during meiotic prophase causes premature loss of arm cohesion in Drosophila oocytes (Perkins *et al*., 2016). Therefore, Sirt1 KD during meiotic prophase could lead to oxidative stress that results in premature loss of arm cohesion. Moreover, SRT1720-mediated activation of Sirt1 may slow reproductive aging and reduce segregation errors by limiting aging-induced oxidative stress in the Drosophila oocyte.

Our results indicate that when Drosophila oocytes undergo aging, the amount of Sirt1 protein associated with oocyte chromosomes does not change but acetylation of H4K16 on oocyte DNA is significantly higher in aged oocytes. The robust H4K16ac signal intensity in *sirt1* null oocytes and aged oocytes is comparable, suggesting that Sirt1 activity is absent in Drosophila diplotene oocytes after the aging regimen. However, if females are fed SRT1720 during the aging regimen, aging-induced loss of Sirt1 activity is prevented and age-dependent chromosome segregation errors are significantly reduced. These results complement the findings in mice that Sirt1 is required to maintain oocyte quality during aging (Di Emidio *et al*., 2014; Iljas *et al*, 2020; Vo *et al*, 2023; Zhang *et al*., 2016) and suggest that therapeutic approaches to maintain Sirt1 activity in aging oocytes may provide an effective strategy to reduce age-dependent segregation errors.

If Sirt1 protein levels do not decrease, why does Sirt1 activity decline with age in Drosophila oocytes? One possibility is that NAD^+^ decreases during aging and becomes limiting as a Sirt1 cofactor. Diminution of NAD^+^ during aging has been reported for several tissues in mice and also in human tissues, although sample sizes in the latter are small (McReynolds *et al*, 2020). Notably, dietary supplementation with a NAD^+^ precursor decreases aneuploidy in chromosome spreads of meiosis II oocytes from aged mice (Miao *et al*, 2020), consistent with Sirt1 activation decreasing age-dependent segregation errors.

While we acknowledge that using Drosophila as a model for aging human oocytes has limitations, we consider this system a valuable tool to better understand mechanisms underlying the maternal age effect. Major strengths include the short generation time, a variety of genetic tools and a simple assay to quantify meiotic segregation errors in oocytes. In addition, our age-dependent NDJ assay (using *mtrm^KG^ smc111* heterozygotes) is sensitized for detection of segregation errors arising from premature loss of cohesion. Although we limit our aging regimen to 4 days for technical reasons, stage 7 & 8 oocytes spend 11-18X more time in diplotene when they arrest and age (Subramanian & Bickel, 2008). Furthermore, only diplotene oocytes are vulnerable to age-dependent NDJ; compared to non-aged oocytes, segregation errors are not elevated in oocytes that arrest and age prior to synaptonemal complex disassembly (Subramanian & Bickel, 2008). One potential limitation of our experiments is that Drosophila females were fed SRT1720 during the entire time-period that their oocytes underwent aging, a protocol that would be challenging to replicate for mammalian oocytes. Further work will be required to determine whether supplementation partway through the aging regimen can still significantly reduce age-dependent segregation errors.

In conclusion, our data demonstrate that preserving Sirt1 function during aging prevents age-dependent chromosome segregation errors in Drosophila oocytes. We hope these findings will inform further exploration of Sirt1 activation as a mechanism to preserve the fidelity of chromosome segregation as oocytes age.

## METHODS

### Reagents and Tools Table

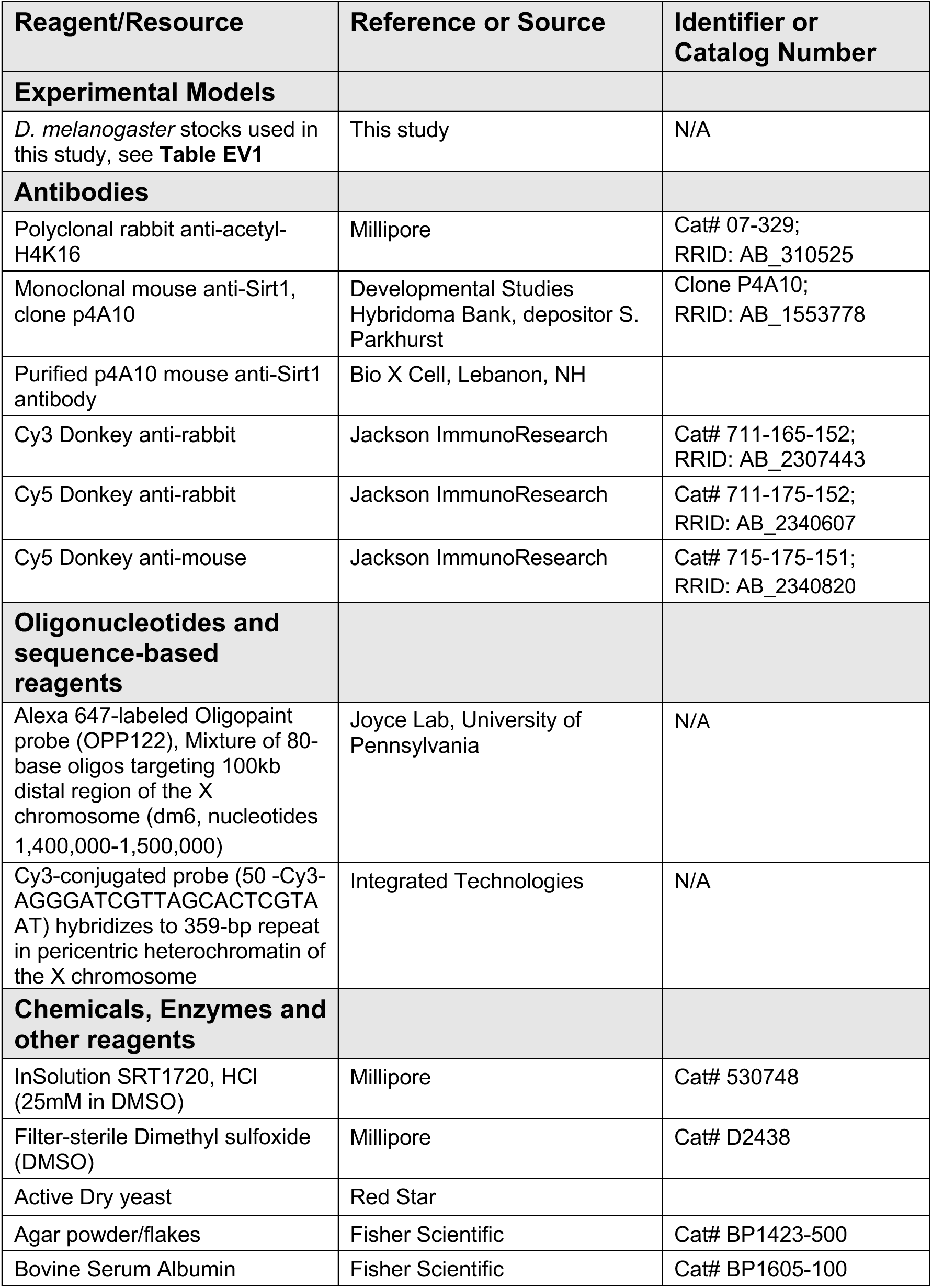
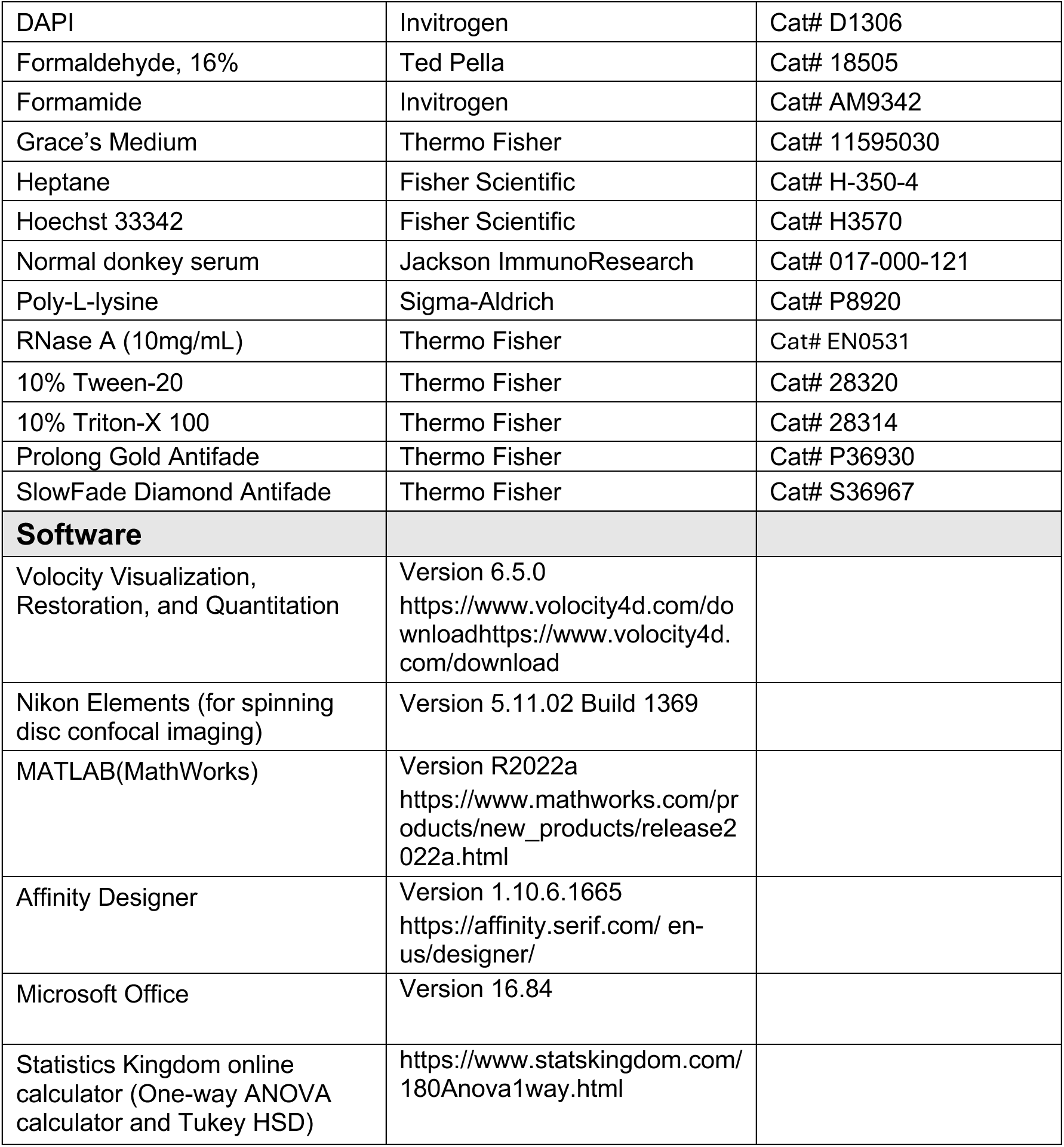

### Methods and Protocols

#### Fly stocks and crosses

Stocks and crosses were raised on standard cornmeal-molasses food and kept at 25°C in a humidified incubator. **Table EV1** provides full genotypes for all stocks utilized in this study as well as Bickel stock numbers which accompany the cross descriptions below.

#### X-chromosome NDJ and Recombinational History assays

To measure chromosome segregation errors in control and Sirt1 KD oocytes, *y sc cv v f car; +; mtrm^KG^*/TM3,Sb (M-835) or *y sc cv v f car; +; mtrm^KG^, matα/TM3,Sb* (M-834) males were crossed to *y* virgins containing a UAS-Sirt1 short hairpin (SH) insertion on the 2^nd^ chromosome (*Sirt1^SH022-B06^*, H-087) or the 3^rd^ chromosome (*Sirt1^SH00806^,* H-084). Non-balancer **Control** (*mtrm^KG^,* no driver ➔ *Sirt1^SH^*) and **KD** (*mtrm^KG^,* matα driver ➔ *Sirt1^SH^*) female progeny were collected as virgins and mated to *X^Y, Bar* (C-200) males in vials containing food and a small amount of dry yeast. For each resulting Sirt1 control and KD genotype, 20 vials of the NDJ cross (8 virgins × 4 males) were set on day 0, the parents cleared on day 7, and the progeny scored daily from day 11 through day 18 Because all the progeny from normal gametes but only half of the progeny from exceptional gametes survive **(Fig EV1C),** % NDJ is calculated using the following formula: [(2(Diplo-X + Nullo-X))/(N + Diplo-X + Nullo-X)]100, where N is the total number of progeny scored. To determine whether Sirt1 KD significantly impacted the fidelity of meiotic chromosome segregation, P values were calculated for each hairpin by comparing control and KD data using the method described in (Zeng *et al*, 2010).

The X-chromosome genotype of the females used for the NDJ assay (*y/ y sc cv v f car)* allowed us to perform an additional test to determine the recombinational history of the missegregating X chromosomes inherited by the Diplo-X progeny (**Fig EV1D**). When scoring the NDJ tests, Diplo-X progeny were collected each day and phenotyped for the X-chromosome visible markers *sc, cv*, *f,* and *car*. Each female was mated to two *y w* (A-062) males and the parents cleared on day 7. Male progeny were scored for *sc, cv*, *f,* and *car* through day 18. By considering the X-chromosome phenotype of the Diplo-X female and her sons, one may deduce the genotype of the two X chromosomes that missegregated, including whether they recombined before missegregation and whether they were sisters (*car^+/+^* or *car^-/-^)* or homologs (*car^+/-^).* The frequency at which recombinant X chromosomes underwent missegregation in each genotype was calculated by dividing the number of Diplo-X females that inherited at least one recombinant chromosome by the total number of progeny scored in the NDJ test and multiplying by 100. Meiosis I and Meiosis II errors were graphed individually and based on whether the Diplo-X female inherited two homologs (Meiosis I) or two sisters (Meiosis II). A two-tailed 2 x 2 *chi^2^* contingency test with Yates’ correction (GraphPad) was used to determine whether the frequency of Diplo-X females with at least one recombinant X chromosome was significantly different between Sirt1 KD and control. P value calculations were performed separately for Diplo-X females that inherited two homologs (Meiosis I) and Diplo-X females that inherited two sisters (Meiosis II).

Not all Diplo-X females were fertile, and a small number of Diplo-X genotypes could not be confidently called. This assay will underestimate the number of recombinant bivalents because only two of the four chromatids will be inherited by the Diplo-X female (and therefore scored) and double crossovers within the large interval between *cv* and *f* will be invisible. In addition, although crossovers are unlikely within the 3.5 cM interval between *car* and heterochromatin, a small number may still occur.

#### FISH to score for cohesion defects

We utilized two X chromosome probes so that we could score for arm and pericentric cohesion defects for the same chromosome assayed in the NDJ and Recombinational History tests. The Alexa 647-labeled OligoPaint probe (OPP122) contains a mixture of 80-mers that recognize a 100-kb distal region on the X chromosome and allows us to assess the state of arm cohesion. The Cy3-labeled pericentromeric probe hybridizes to an 11 Mb stretch of satellite DNA near the centromere of the X chromosome. Unfortunately, the large size of this target may limit our detection of cohesion defects near the X chromosome centromere (Haseeb *et al*., 2024a; Haseeb *et al*., 2024b).

To generate control & KD genotypes, *y w* (A-062) or *matα* (T-273) males were crossed to *Sirt1^SH022-B06^* virgins (H-213). 20-25 young female progeny were held with 10 males in a vial with food and dry yeast for 3 days before dissection.

Below is a brief description of the steps used in FISH detection of sister chromatid cohesion defects in mature Drosophila oocytes (stages 13-14). Additional details, especially regarding removal of chorions and vitelline membranes and tips for oocyte handling, may be found in (Perkins & Bickel, 2017). A detailed protocol is available upon request.

In a shallow dissecting dish, 15-20 sets of ovaries were dissected within a 10 min window in 1X Modified Robb’s buffer (55 mM potassium acetate, 40 mM sodium acetate, 100 mM sucrose, 10 mM glucose, 1.2 mM magnesium chloride, 1.0 calcium chloride, 100 mM Hepes, pH 7.4) and transferred to a 1.5mL microfuge tube. After addition of 500 µL prewarmed (37°C) fixative (4% formaldehyde, 100 mM sodium cacodylate, 100 mM sucrose, 40 mM sodium acetate, 40 mM sodium acetate, 10 mM EGTA) and 500µL of heptane (RT), the tube was mixed vigorously and ovaries fixed for 6 min on a nutating mixer. Following removal of fixative, ovaries were washed three times with 500 µL of 1X PBSBTx (1X PBS / 0.5% BSA / 0.1% Triton X-10) by inverting tube 2-3 times and allowing ovaries to settle. Fixed ovaries were transferred to a shallow dissecting dish containing PBSBTx and pipetted up and down using a BSA-coated gel-loading tip to help dissociate the later stages. This mixture was rolled between the frosted regions of two slides to manually remove the chorion and vitelline membrane of mature oocytes and transferred to a 15mL conical tube for three rounds of settling (3mL of PBSBTx) to remove younger stages and debris which settle less quickly. Mature oocytes were transferred to a 500µL microfuge tube and stored in PBSBTx overnight at 4°C. The next morning, oocytes were rinsed once with 500µL PBSTx (1X PBS containing 1% Triton X-100) and incubated on a nutating mixer for 2 hours at room temperature (RT) in PBSTx containing 100µg/mL RNAse). Following three rinses in 2X SSCT (0.3M sodium chloride, 30mM sodium citrate with 0.1% Tween 20), oocytes were washed three times 10 min in 2X SSCT, followed by a 10 min wash in 2X SSCT containing 20% formamide, another with 2X SSCT / 40% formamide and a final wash in 2X SSCT / 50% formamide, all at RT on a nutating mixer. Oocytes were incubated in 2X SSCT + 50% formamide with rotation for 2 hours at 37°C before transfer to 200µL PCR tubes and a pre-denaturation program in a thermocycler (5min @ 37C, 3 min @ 92C, 20 min @ 60C, Hold @ 37C). Probes in 50µL of 1X hybridization buffer (3X SSC, 50% formamide, 10% dextran sulfate) were allowed to hybridize overnight after denaturation (37°C for 5 min, 92 °C for 3 min, Hold @ 37 °C). The Cy3-labeled pericentromeric probe was used at 1 ng/µL and the Alexa 647-labeled Oligopaint probe (OPP122) was used at 0.50 pmol/µL. On day 3, oocytes were transferred to a pre-warmed 500µL microfuge tube and taken through a series of 500µL washes at 37°C on a rotator: three 20 min washes in 2X SSCT / 50% formamide, three 10 min washes in 2X SSCT / 50% formamide. Washes continued at RT on a nutating mixer: one 10 min wash in 2X SSCT / 40% formamide, one 10-min wash in 2X SSCT / 20% formamide and one 10 min wash in 2X SSCT. Following a 30 min incubation with DAPI (1 µg/mL) in 2X SSCT (protected from light on a nutating mixer), oocytes were rinsed 3 times with 2X SSCT followed by two 10 min washes in 2X SSCT. Oocytes were rinsed with 1X PBS / 0.01% Tween-20 and left in that solution for mounting onto poly-L-lysine coated 18 mm #1.5 coverslips with 25µL of Prolong Gold mounting medium. Slides were allowed to cure in the dark for at least 21 days before imaging.

#### Sirt1 and H4K16ac Immunostaining

*matα* (T-273) or *y w* (A-062) males were crossed to *Sirt1^SH022-B06^*virgins (H-213) to produce Sirt1 control and KD ovaries/oocytes. To generate *sirt1* null oocytes, *sirt1^5.26^*males were crossed to *sirt1^4.5^* females and ovaries of female progeny were examined. Each of these *sirt1* deletion alleles removes >750 nucleotides of the Sirt1 coding sequence but leaves the coding sequence of the adjacent *DnaJ-H* gene intact (Newman *et al*, 2002). For the above genotypes, newly eclosed females were held with males for 2-3 days in vials with food and a small amount of dry yeast before ovary dissection

For immunostaining aged and non-aged ovarioles, *mtrm^KG^ smc1Δ* / TM3 males (M-822) were crossed to *y w* females (A-062). Non-balancer virgins from this cross were collected during an 8-12 hour interval, held overnight in vials with food and a small amount of dry yeast, and subjected to the 4-day aging regimen (**Fig EV2**, and described in more detail below). Ovary dissections were performed immediately following the aging regimen.

For each genotype or treatment condition, 6 sets of ovaries were dissected in Grace’s medium and ovarioles were gently splayed (early stages) using a fine tungsten needle. Fixation was performed in a deep-well glass dish with gentle rotation on a shaker for 20 min at RT in 400µL of 1X PBS containing 2% formaldehyde. All subsequent incubations and washes were performed in a glass dish at RT with gentle rotation on a shaker. Following fixation, ovaries were rinsed three times with 400µL of 1X PBS / 0.2% Triton X-100 and permeabilized by performing two 15-min incubations in 400µL of 1X PBS containing 0.5% Triton X-100. After three rinses with 400µL of 1X PBS / 0.2% Tween-20, ovaries were blocked for one hour in 400µL of blocking buffer (1X PBS / 0.2% Tween-20 / 0.5% BSA / 5% Donkey Serum). Ovaries were incubated overnight in a humidified box in 200µL of 1X PBS / 0.01% Tween-20 / 0.5% BSA containing primary antibody. The next morning, 400µL of 1X PBS / 0.2% Tween-20 was used for each of three rinses followed by three 20 min washes. Ovaries were incubated for one hour, protected from light, in 200µL in 1X PBS / 0.01% Tween-20 / 0.5% BSA containing secondary antibody. Following three rinses and a 20 min wash with 400µL of 1X PBS / 0.2% Tween-20, ovaries were incubated 20 min in 1X PBS containing Hoechst (2.0µg/mL), and 20 min in 1X PBS / 0.01% Tween-20. After separation with fine tungsten needles, ovarioles were transferred onto 18mm poly-L-lysine coated #1.5 coverslips, excess liquid removed and a slide with mounting medium lowered onto the coverslip.

Either SlowFade Diamond (Fig 3 & 4) or ProLong Gold (Fig 5) was used for mounting. SlowFade Diamond provided the highest signal to noise ratio for anti-Sirt1 staining, but the signal did degrade over the course of a week, so all images were captured within three days of mounting. Coverslips were sealed with nail polish immediately upon SlowFade mounting and slides stored flat and protected from light at 4°C. For ProLong Gold, slides were allowed to cure in the dark for at least 14 days before application of nail polish and imaging.

Purified anti-Sirt1 antibody (∼52mg at 3.7 mg/mL) was produced by Bio X Cell (Lebanon, NH) using the mouse hybridoma cell line P4A10 that we obtained from the Developmental Studies Hybridoma Bank (DSHB). Mouse anti-Sirt1 antibody was used at 1µg/mL, and rabbit anti-H4K16ac (Millipore Cat # 07-329) was used at 1:100. Donkey secondary antibodies (Jackson ImmunoResearch) were used at a final dilution of 1:400. For Figs 3 & 4, Cy3 anti-rabbit (711-165-152) was used to detect H4K16ac and Cy5 anti-mouse (715-175-151) was used to detect Sirt1. For Fig 5, Cy5 anti-rabbit (711-175-152) was used to detect H4K16ac. For Fig 3, samples were incubated simultaneously with both primary antibodies. For Fig 4, samples were incubated with either anti-Sirt1 or anti-H4K16ac.

#### Aging regimen

For all experiments that compared aged and non-aged oocytes, *mtrm^KG^ smc1Δ / TM3* males (M-822) were crossed to *y w* virgins (A-062) to generate *mtrm^KG^ smc1Δ / +* virgins that were collected during an 8-12 hour window and held in vials overnight with food and a small amount of dry yeast. At approximately 2 pm the following day, virgins were divided equally and placed in laying bottles (**Fig S2).** In one laying bottle, virgins were held in the absence of males. Because egg-laying is suppressed in the absence of mating, oogenesis halts in these females and stages 8 and earlier arrest and “age” in the female (Subramanian & Bickel, 2008). Females placed into a laying bottle with *X^Y, Bar* (C-200) males will lay fertilized eggs continuously during the aging regimen and provide the source for “non-aged” oocytes.

Every 24 hours, flies in laying bottles were provided with a fresh 60 mm petri plate containing 5% glucose / 2% agar as well as a smear (∼ 1 cm diameter) of newly prepared yeast paste (0.6 g in 1mL sterile ultrapure water) on the agar surface. At the end of each 24-hr interval, the surface of each agar plate was photographed to document egg laying (or lack thereof) for each laying bottle. Upon completion of the 4-day aging regimen, flies were transferred from the laying bottles back into vials and used for cytology or NDJ experiments. We limit our aging regimen to a 4-day time frame because virgins lay an increased number of unfertilized eggs after 4 days.

#### SRT1720 supplementation

25mM SRT1720 in DMSO (Millipore, Cat # 530748) was aliquoted upon arrival, stored at -80°C, and a fresh aliquot used for each experiment. On the first day of an aging regimen, one aliquot of 25mM SRT1720 was diluted to 10mM using a thawed aliquot of DMSO (Millipore, Cat # D2438), also stored at -80°C. Both the 10mM SRT1720 solution and the remaining thawed DMSO were kept at 4°C during the 4-day aging regimen. On each day of the aging regimen, a fresh 2mM SRT1720 solution was prepared using sterile ultrapure water. 5µL of 2mM SRT1720 was added to 995µL of sterile ultrapure water to achieve the final concentration of 10uM SRT1720, and 0.6g of yeast was added to this solution to make yeast paste.

For the control group, we matched the concentration of DMSO in yeast paste (0.1%) to that for the SRT1720 treatment. On each day of the aging regimen, 5µL of 100% DMSO was added to 20µL of sterile ultrapure water to generate a 20% DMSO solution. 1mL of 0.1% DMSO was prepared by adding 5µL of 20% DMSO to 995µL of sterile ultrapure water and 0.6g of yeast was added to this solution to generate the yeast paste.

#### Age-Dependent NDJ

To compare the frequency of segregation errors in aged and non-aged oocytes, *mtrm^KG^ smc1Δ / +* females were removed from laying bottles at the end of the 4-day aging regimen and used to set up NDJ crosses in vials with *X^Y, Bar* (C-200) males. A smear of wet yeast (with no SRT1720 or DMSO addition) was applied to the wall of each vial. In most cases, 10 vials (4 females +3 males) were set for each condition (aged + DMSO, non-aged + DMSO, aged + SRT1720, non-aged + SRT1720. Parents were removed after 48 hours and progeny scored through day 18.

Data from the three individual experiments are reported in **Table EV2.** The method described in (Zeng *et al*., 2010) was used to determine whether aged oocytes exhibited a significant increase in NDJ compared to non-aged oocytes when mothers were fed DMSO or SRT1720. **Fig 5D** graphs the means from the three independent experiments. A one-way ANOVA (Statistics Kingdom), using the NDJ values from all three replicates, indicated that there was a significant difference between at least two groups (F(3,8) = [26.8], P = 0.00016). To determine whether NDJ values differed significantly between two specific conditions, a Tukey’s HSD test (Statistics Kingdom) for multiple comparisons was performed and P values for pair-wise comparisons are presented in **Fig 5D**.

#### Image acquisition and processing

All images were acquired with an Andor Spinning Disk confocal (50µm pinhole) using Nikon Elements (5.11.02 Build 1369) to control a Nikon Eclipse Ti inverted microscope, ASI MS-2000 motorized piezo stage, Zyla 4.2-megapixel sCMOS camera and three lasers (405, 561 and 637 nm). All image acquisition utilized 4X frame averaging. For all Z stack imaging, a complete Z stack was captured for each fluor sequentially, moving from longest to shortest wavelength.

For FISH imaging (ROI: 512 x 512), a CFI 100x oil Plan Apo DIC objective (NA 1.45) was used to capture a 4 µm Z series with 0.1 µm steps. For Sirt1 and H4K16ac imaging, a Nikon CFI 40X Plan Apo oil objective (NA 1.4) was used with the following workflow. First, the slide was scanned for ovarioles and the XY position recorded for presumptive stage 7 and stage 8 egg chambers. For each egg chamber imaged, a Z series (0.5um step, 2 um total) was acquired for a small ROI (256 x 256) that contained the oocyte DNA. These images are shown in Figs 3-5. Next, a single optical section was acquired for the entire egg chamber, focusing on a plane that included the oocyte DNA. These images were used for **Fig EV3** Lastly, a full-field (2048 x 2048) Z series (0.5um step, 5um total) was captured that included as much of the ovariole as possible and this image stack was used to confirm the developmental stage of the egg chamber that was imaged in the preceding steps. Staging was based on size and morphological criteria (King, 1970; Mahowald & Kambysellis, 1980; Spradling, 1993).

For all comparisons of genotypes and/or conditions, acquisition and image processing parameters were identical for all Sirt1 images and separately for all H4K16ac images. In addition, when Z series are presented, the same number of optical sections are included for each of the images shown in a series.

### Quantification Methods

#### Scoring cohesion defects

A MATLAB (R2022a) script was used to randomize and rename FISH images so that all FISH scoring was performed blind to sample identity. Image stacks were batch processed for deconvolution (Volocity Restoration, v6.5.0) using identical parameters for all datasets. Cohesion defects were scored (Volocity Visualization, v6.5.0) by manually scrolling in all three dimensions to visualize/count the number of Cy3 and Alexa 547) probe spots on the oocyte DNA. Two spots were scored as separate if the distance between them was greater than one half the diameter of the smaller spot AND there was no evidence of a thread connecting them. Following scoring of both arm and pericentric defects, sample ID was revealed and the percentage of oocytes with cohesion defects calculated. Only arm defects were graphed in **Fig 2** because no pericentric defects were detected in Sirt1 KD or control oocytes. A two-tailed Fisher’s exact test (Graphpad) was utilized to determine whether the percentage of oocytes with arm defects were significantly different in two genotypes.

#### Quantification of anti-Sirt1 and anti-H4K16ac signal on oocyte DNA

Volocity Quantitation (v6.5.0) was used for all image quantification steps. Quantification of Anti-Sirt1 and/or anti-H4K16ac signal intensity on oocyte DNA was performed using the short Z series (0.5um step, 2 um total) acquired as a 256 x 256 ROI (described above). The image was cropped such that the only DNA in the field was oocyte DNA (405 channel). Threshholding was used to identify the voxels containing oocyte DNA (405 nm channel). For every voxel within the DNA volume, Volocity Quantification provided the signal intensity (0-4095) for anti-Sirt1 (637nm channel) and anti-H4K16ac (561nm channel).

In **Figs 3-5**, box and whiskers plots plot the average intensity of Sirt1 or H4K16ac on DNA for each oocyte. In all box and whiskers plots shown, the average intensity is indicated with an “X”, the median and quartiles are depicted by horizontal lines and outliers are included as dots. Outliers are shown as solid dots and correspond to values that lie more than 1.5 * IQR (interquartile range) above the 75% or below the 25% quartile.

A two tailed unpaired t-test (Microsoft Excel) was used to calculate whether the difference between genotypes and/or conditions were significant. P values are provided on the graphs in Figs 3-5 and the number of oocytes scored is shown for each genotype.

## DATA AVAILABILITY

This study includes no data deposited in external repositories.

## AUTHOR CONTRIBUTIONS

**Zihan Meng:** Conceptualization, Investigation, Formal analysis, Visualization, Writing – original draft, Writing – review and editing. **Nicholas G. Norwitz:** Conceptualization, Investigation, Formal analysis, Writing – review and editing. **Sharon E. Bickel:** Funding acquisition, Supervision, Project administration, Resources, Conceptualization, Formal analysis, Visualization, Writing – original draft, Writing – review and editing.

## ACKNOWLEDGEMENTS

All Microscopy was performed in the Dartmouth Life Sciences Light Microscopy Core Facility. We thank Ann Lavanway for microscopy assistance and Britton Johnson for preparation of fly food. We are grateful to Soni Lacefield, Muhammad Abdul Haseeb and Diana Hilpert for comments on the manuscript. We thank the Bloomington Drosophila Stock Center (NIH P40OD018537) for providing fly stocks and the Transgenic RNAi Project (NIH R24OD030002) for producing *Sirt1^SH^* stocks. The p4A10 anti-Sirt1 hybridoma cell line, generated by Susan Parkhurst (Fred Hutchinson Cancer Research Center), was obtained from the Developmental Studies Hybridoma Bank, created by NICHD of the NIH and maintained by the Department of Biology at The University of Iowa.

## SUPPLEMENTAL INFORMATION

**Table EV1.**
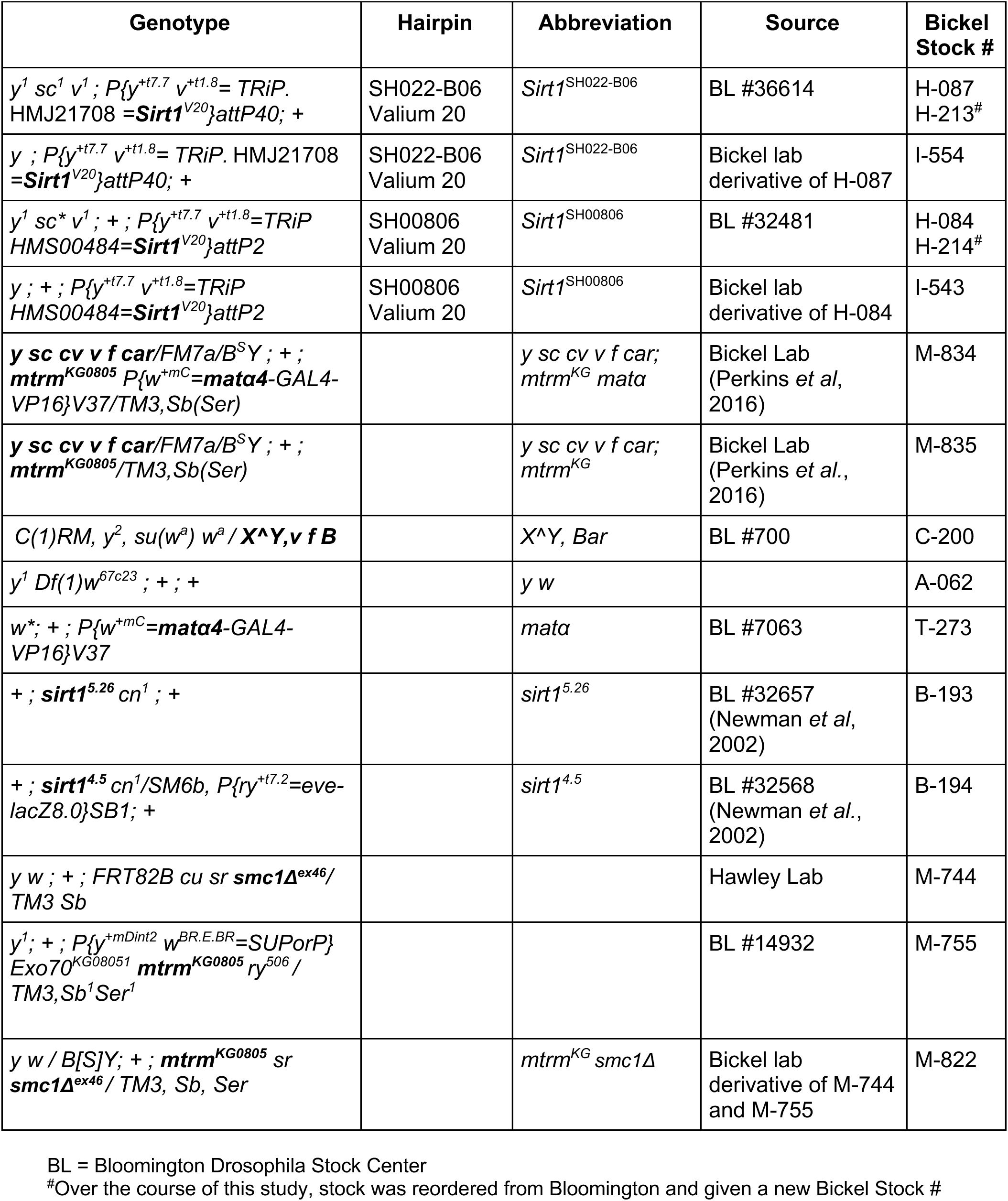
Fly Stocks and Genotypes.

**Table EV2.** Age-dependent NDJ is suppressed by feeding mothers SRT1720.

| Condition | % NDJ | % Diplo (#) | % Nullo (#) | N (fert) | P-value |
| --- | --- | --- | --- | --- | --- |
| **Replicate 1** |  |  |  |  |  |
| Aged, DMSO | 15.26 | 7.07 (57) | 8.19 (66) | 1489 (37.2) | 7.8 x 10<sup>-6</sup> |
| Non-Aged, DMSO | 7.83 | 4.18 (31) | 3.64 (27) | 1424 (35.6) |  |
| Aged, SRT1720 | 12.79 | 6.01 (46) | 6.79 (52) | 1434 (35.9) | 0.126 |
| Non-Aged, SRT 1720 | 10.19 | 4.13 (30) | 6.06 (44) | 1378 (34.5) |  |
| **Replicate 2** |  |  |  |  |  |
| Aged, DMSO | 15.12 | 8.21 (38) | 6.91 (32) | 856 (23.8) | 5.35 x 10<sup>-4</sup> |
| Non-Aged, DMSO | 8.02 | 4.16 (27) | 3.86 (25) | 1245 (34.6) |  |
| Aged, SRT1720 | 11.43 | 5.87 (37) | 5.56 (35) | 1188 (29.7) | 0.111 |
| Non-Aged, SRT 1720 | 8.71 | 5.37 (37) | 3.34 (23) | 1318 (33.0) |  |
| **Replicate 3** |  |  |  |  |  |
| Aged, DMSO | 15.42 | 8.57 (35) | 6.85 (28) | 754 (18.9) | 2.48 x 10<sup>-3</sup> |
| Non-Aged, DMSO | 8.51 | 3.40 (16) | 5.11 (24) | 900 (22.5) |  |
| Aged, SRT1720 | 9.56 | 3.74 (18) | 5.82 (28) | 916 (22.9) | 0.232 |
| Non-Aged, SRT 1720 | 7.43 | 2.18 (12) | 5.52 (29) | 1063 (26.5) |  |

In all three experiments, the genotype of mothers was: *y w/y w; + : mtrm<sup>KG</sup> smc1Δ / +*.

---

**Figure EV1.**
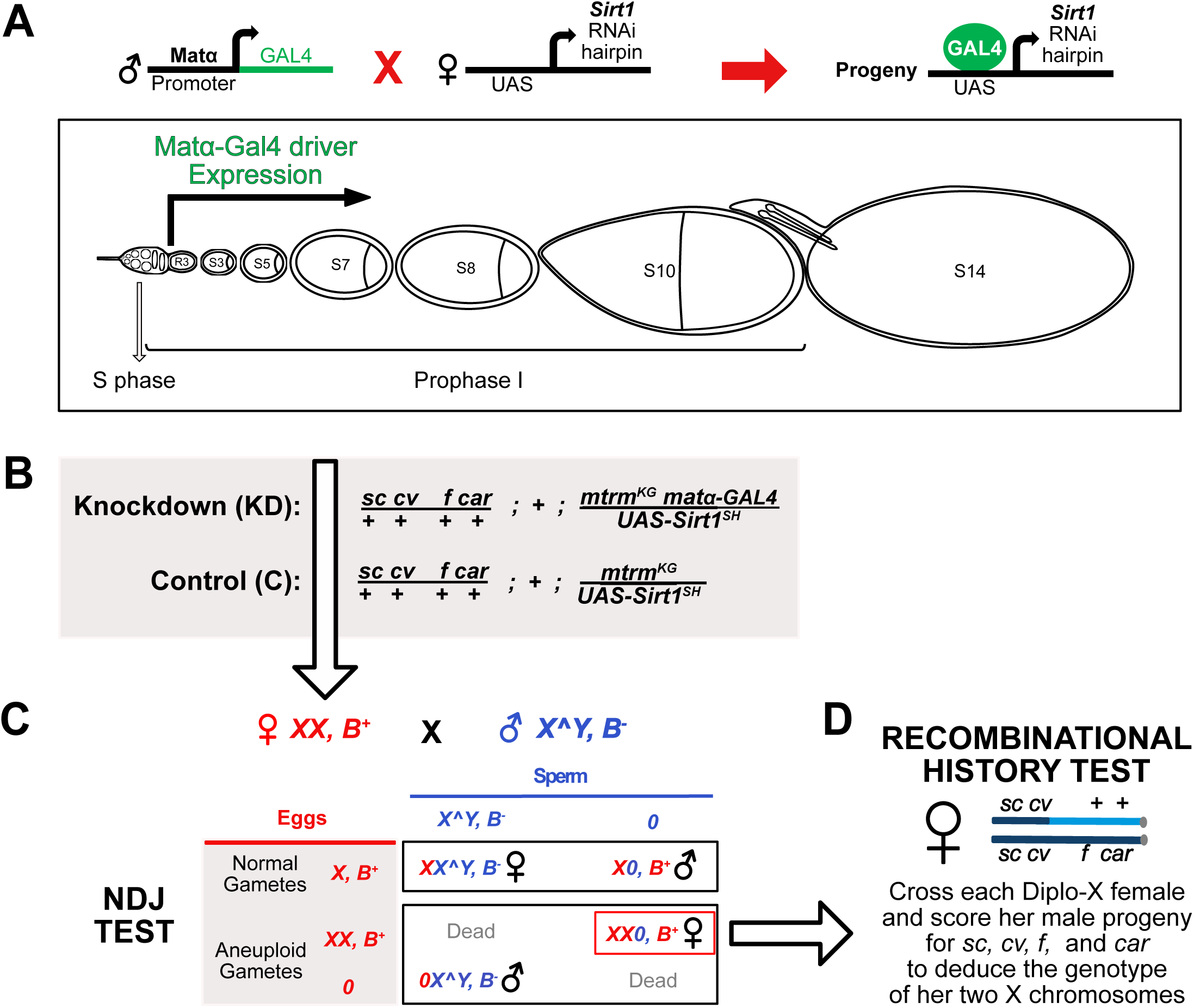
Genetic tools and tests to assay for meiotic segregation errors. **A.** Gal4/UAS strategy to knock down Sirt1 in the Drosophila female germline using the matα-Gal4 driver. Expression of the matα driver does not begin until Region 3 (R3) or stage 2 of oogenesis, at least two days after meiotic cohesion is established during S phase. Note that a single ovariole does not contain all stages concurrently. **B.** Genotypes are shown for Sirt1 Knockdown and Control females used in the NDJ test with the *Sirt1^SH^* transgene on the 3^rd^ chromosome. Note that both genotypes are heterozygous for the *mtrm^KG^* allele which weakens arm cohesion and also makes it more likely that premature loss of cohesion will result in segregation errors in our NDJ assay. Females are also heterozygous for four visible makers on the X chromosome (*sc, cv, f & car*) which permit the recombinational history analysis following the NDJ test. **C.** In the NDJ test shown, X-chromosome segregation errors in oocytes can be quantified because progeny arising from normal and aneuploid gametes survive and can be counted. Sirt1 KD or C females are mated to attached *X^Y*, *B* males and progeny scored based on sex and the dominant marker *Bar*. Only half of the Diplo-X and Nullo-X eggs will be viable when fertilized. **D.** Diplo-X females inherit both X chromosomes from their mother due to missegregation in the oocyte. By mating Diplo-X females and scoring their male progeny for the X chromosome markers *sc, cv, f* and *car*, we can determine whether a recombinant bivalent underwent missegregation and, based on *car*, whether the two inherited X chromosomes were homologs (Meiosis I error) or sisters (Meiosis II error). In the example shown here, the Diplo-X female inherited one recombinant and one non-recombinant X chromosome and heterozygosity for the mutant *car* allele indicates that the two X chromosomes are homologs.

**Figure EV2.**
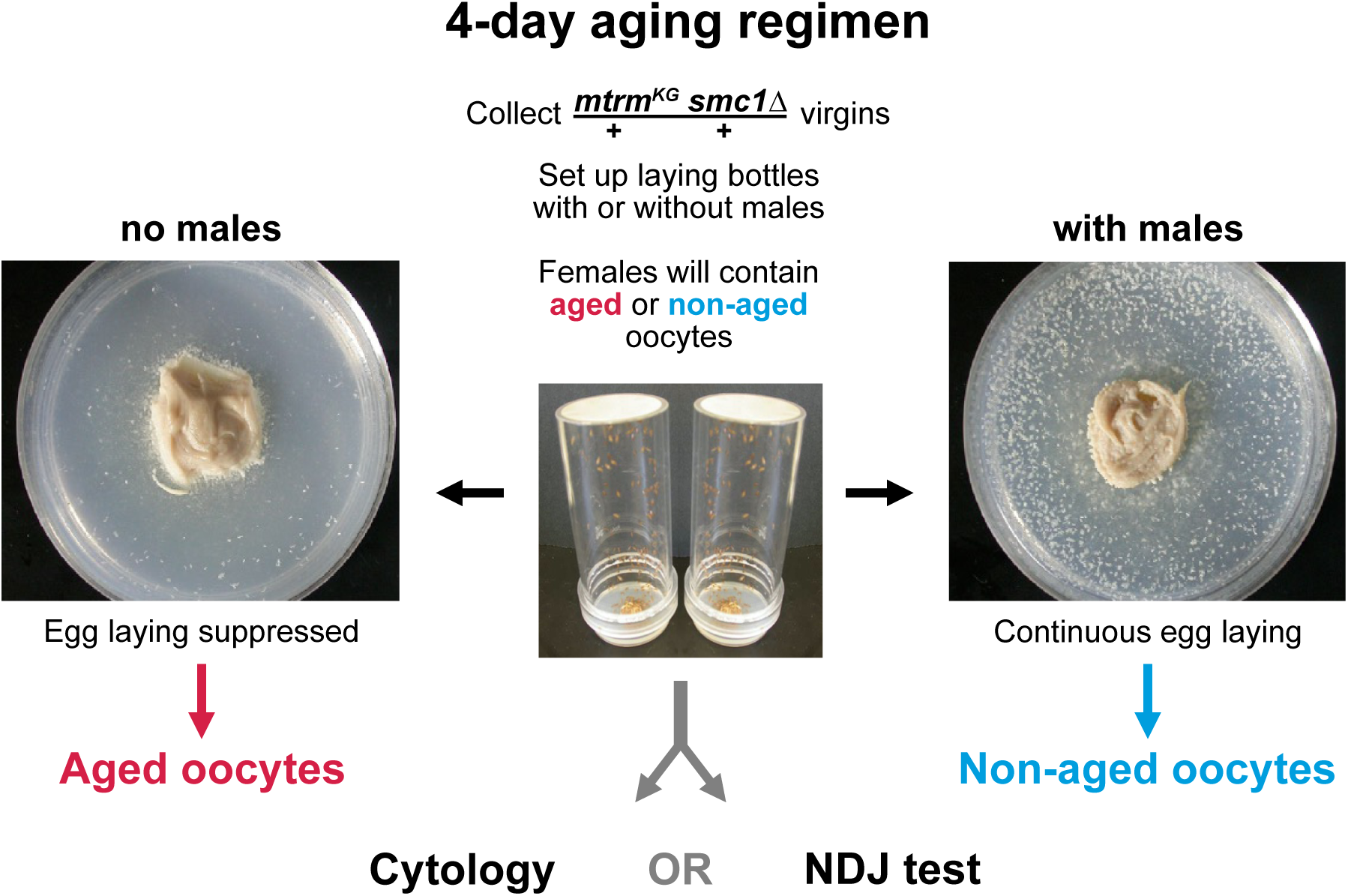
Experimental procedure to generate and compare aged and non-aged oocytes. *mtrm^KG^ smc1Δ / +* virgins were used for all aging experiments. In this genotype, cohesion is weakened but not abolished and disruption of the achiasmate segregation system allows bivalents that lose their chiasma (due to loss of arm cohesion) to missegregate. Virgins are placed in plastic laying bottles with a glucose/agar plate and yeast paste and *X^Y, B* males are omitted (left, aged) or added (right, non-aged). **Left:** In the absence of mating, egg laying is suppressed and most oogenesis stages halt progression. On the agar plate shown, very few unfertilized eggs (white spots on agar surface) have been laid during the 24-hr interval. When oogenesis halts, oocytes arrest and age at a particular stage. Drosophila oocytes are most vulnerable to aging-induced segregation errors when they arrest and age in diplotene. **Right:** Oogenesis is stimulated in females that have mated, and many fertilized eggs are laid within a 24-hr period. Because oogenesis is continuous in these females, they produce non-aged oocytes.

**Figure EV3.**
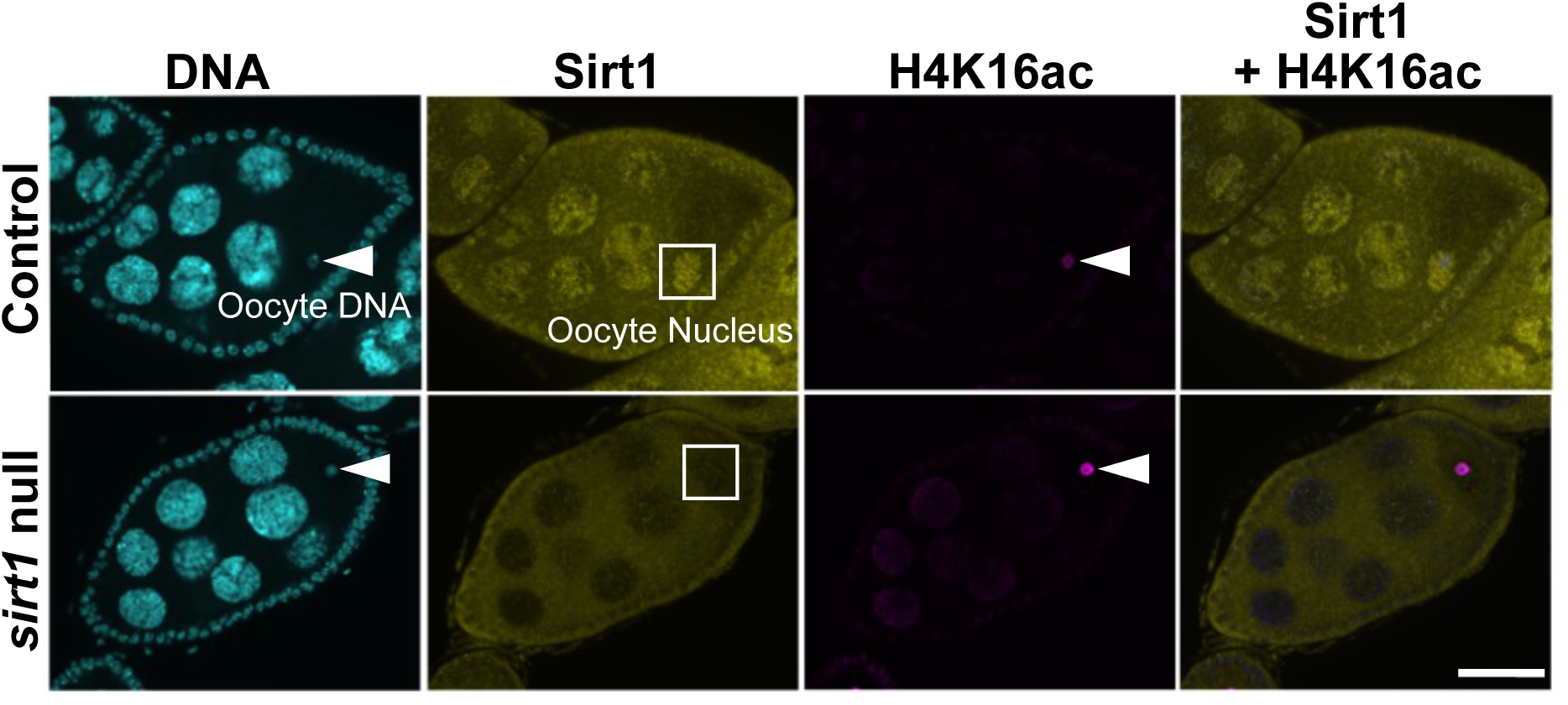
Sirt1 and H4K16ac immunolocalization in stage 7 egg chambers. A single confocal optical section is shown for Sirt1 and H4K16ac immunostaining in stage 7 egg chambers of Control (no driver **→** short hairpin) and *sirt1* null (*sirt1^4.5^/sirt1^5.26^*) genotypes. DNA signal is cyan, Sirt1 staining is yellow, and the H4K16 acetylation signal is magenta. Scale bar, 20 µm. The arrowheads point to the oocyte DNA. A white square surrounds each oocyte nucleus. In Control egg chambers, Sirt1 immunostaining is enriched in the nuclei of nurse cells, follicle cells and the oocyte but this pronounced nuclear Sirt1 signal is absent in *sirt1* null egg chambers. H4K16ac signal is faintly visible on the chromosomes of Control oocytes. In contrast, the H4K16ac signal on oocyte DNA is significantly stronger in *sirt1* null oocytes, consistent with lack of Sirt1 deacetylase activity in this genotype.

## REFERENCES

Alam F, Syed H, Amjad S, Baig M, Khan TA, Rehman R (2021) Interplay between oxidative stress, SIRT1, reproductive and metabolic functions. Curr Res Physiol 4: 119-124

Bickel SE, Orr-Weaver T, Balicky EM (2002) The sister-chromatid cohesion protein ORD is required for chiasma maintenance in Drosophila oocytes. Curr Biol 12: 925–929

Bonner AM, Hughes SE, Hawley RS (2020) Regulation of Polo Kinase by Matrimony Is Required for Cohesin Maintenance during Drosophila melanogaster Female Meiosis. Curr Biol 30: 715–722 e713

Buonomo SB, Clyne RK, Fuchs J, Loidl J, Uhlmann F, Nasmyth K (2000) Disjunction of homologous chromosomes in meiosis I depends on proteolytic cleavage of the meiotic cohesin Rec8 by separin. Cell 103: 387–398

Chang HC, Guarente L (2014) SIRT1 and other sirtuins in metabolism. Trends Endocrinol Metab 25: 138–145

Charalambous C, Webster A, Schuh M (2023) Aneuploidy in mammalian oocytes and the impact of maternal ageing. Nat Rev Mol Cell Biol 24: 27–44

Chen C, Zhou M, Ge Y, Wang X (2020) SIRT1 and aging related signaling pathways. Mech Ageing Dev 187: 111215

Chen Y, Zhao W, Yang JS, Cheng Z, Luo H, Lu Z, Tan M, Gu W, Zhao Y (2012) Quantitative acetylome analysis reveals the roles of SIRT1 in regulating diverse substrates and cellular pathways. Mol Cell Proteomics 11: 1048–1062

Chen YF, Chou CC, Gartenberg MR (2016) Determinants of Sir2-Mediated, Silent Chromatin Cohesion. Mol Cell Biol 36: 2039–2050

Dai H, Sinclair DA, Ellis JL, Steegborn C (2018) Sirtuin activators and inhibitors: Promises, achievements, and challenges. Pharmacol Ther 188: 140–154

Dernburg AF, Sedat JW, Hawley RS (1996) Direct evidence of a role for heterochromatin in meiotic chromosome segregation. Cell 86: 135–146

Di Emidio G, Falone S, Vitti M, D’Alessandro AM, Vento M, Di Pietro C, Amicarelli F, Tatone C (2014) SIRT1 signalling protects mouse oocytes against oxidative stress and is deregulated during aging. Hum Reprod 29: 2006–2017

Gong H, Pang J, Han Y, Dai Y, Dai D, Cai J, Zhang TM (2014) Age-dependent tissue expression patterns of Sirt1 in senescence-accelerated mice. Mol Med Rep 10: 3296–3302

Grabowska W, Sikora E, Bielak-Zmijewska A (2017) Sirtuins, a promising target in slowing down the ageing process. Biogerontology 18: 447–476

Greaney J, Wei Z, Homer H (2018) Regulation of chromosome segregation in oocytes and the cellular basis for female meiotic errors. Hum Reprod Update 24: 135–161

Harris D, Orme C, Kramer J, Namba L, Champion M, Palladino MJ, Natzle J, Hawley RS (2003) A deficiency screen of the major autosomes identifies a gene (matrimony) that is haplo-insufficient for achiasmate segregation in Drosophila oocytes. Genetics 165: 637–652

Haseeb MA, Bernys AC, Dickert EE, Bickel SE (2024a) An RNAi screen to identify proteins required for cohesion rejuvenation during meiotic prophase in Drosophila oocytes. G3 (Bethesda) 14

Haseeb MA, Weng KA, Bickel SE (2024b) Chromatin-associated cohesin turns over extensively and forms new cohesive linkages in Drosophila oocytes during meiotic prophase. Curr Biol 34: 2868–2879 e2866

Hawley RS, Irick H, Zitron AE, Haddox DA, Lohe A, New C, Whitley MD, Arbel T, Jang J, McKim K et al (1992) There are two mechanisms of achiasmate segregation in Drosophila, one of which requires heterochromatic homology. Dev Genet 13: 440–467

Hodges CA, Revenkova E, Jessberger R, Hassold TJ, Hunt PA (2005) SMC1beta-deficient female mice provide evidence that cohesins are a missing link in age-related nondisjunction. Nat Genet 37: 1351–1355

Hubbard BP, Sinclair DA (2014) Small molecule SIRT1 activators for the treatment of aging and age-related diseases. Trends Pharmacol Sci 35: 146–154

Iljas JD, Wei Z, Homer HA (2020) Sirt1 sustains female fertility by slowing age-related decline in oocyte quality required for post-fertilization embryo development. Aging Cell 19: e13204

Januschke J, Gervais L, Dass S, Kaltschmidt JA, Lopez-Schier H, St Johnston D, Brand AH, Roth S, Guichet A (2002) Polar transport in the Drosophila oocyte requires Dynein and Kinesin I cooperation. Curr Biol 12: 1971–1981

Karpen GH, Le MH, Le H (1996) Centric heterochromatin and the efficiency of achiasmate disjunction in *Drosophila* female meiosis. Science 273: 118–122

King RC (1970) Ovarian development in Drosophila melanogaster. Academic Press, New York, NY

Li H (2014) Sirtuin 1 (SIRT1) and Oxidative Stress. In: Systems Biology of Free Radicals and Antioxidants, Laher I. (ed.) pp. 417-435. Springer Berlin Heidelberg: Berlin, Heidelberg

Lopez-Otin C, Blasco MA, Partridge L, Serrano M, Kroemer G (2013) The hallmarks of aging. Cell 153: 1194–1217

Ma R, Zhang Y, Zhang L, Han J, Rui R (2015) Sirt1 protects pig oocyte against in vitro aging. Anim Sci J 86: 826–832

Mahowald A, Kambysellis M (1980) Oogenesis. In: The Genetics and Biology of Drosophila, Ashburner M., Wright T. (eds.) pp. 141-224. Academic Press: New York

McBurney MW, Clark-Knowles KV, Caron AZ, Gray DA (2013) SIRT1 is a Highly Networked Protein That Mediates the Adaptation to Chronic Physiological Stress. Genes Cancer 4: 125–134

McReynolds MR, Chellappa K, Baur JA (2020) Age-related NAD(+) decline. Exp Gerontol 134: 110888

Miao Y, Cui Z, Gao Q, Rui R, Xiong B (2020) Nicotinamide Mononucleotide Supplementation Reverses the Declining Quality of Maternally Aged Oocytes. Cell Rep 32: 107987

Mihalas BP, Marston AL, Wu LE, Gilchrist RB (2024) Reproductive Ageing: Metabolic contribution to age-related chromosome missegregation in mammalian oocytes. Reproduction 168

Milne JC, Lambert PD, Schenk S, Carney DP, Smith JJ, Gagne DJ, Jin L, Boss O, Perni RB, Vu CB et al (2007) Small molecule activators of SIRT1 as therapeutics for the treatment of type 2 diabetes. Nature 450: 712–716

Minor RK, Baur JA, Gomes AP, Ward TM, Csiszar A, Mercken EM, Abdelmohsen K, Shin YK, Canto C, Scheibye-Knudsen M et al (2011) SRT1720 improves survival and healthspan of obese mice. Sci Rep 1: 70

Mitchell SJ, Martin-Montalvo A, Mercken EM, Palacios HH, Ward TM, Abulwerdi G, Minor RK, Vlasuk GP, Ellis JL, Sinclair DA et al (2014) The SIRT1 activator SRT1720 extends lifespan and improves health of mice fed a standard diet. Cell Rep 6: 836–843

Newman BL, Lundblad JR, Chen Y, Smolik SM (2002) A Drosophila homologue of Sir2 modifies position-effect variegation but does not affect life span. Genetics 162: 1675–1685

Park SU, Walsh L, Berkowitz KM (2021) Mechanisms of ovarian aging. Reproduction 162: R19–R33

Perkins AT, Bickel SE (2017) Using Fluorescence In Situ Hybridization (FISH) to Monitor the State of Arm Cohesion in Prometaphase and Metaphase I Drosophila Oocytes. J Vis Exp

Perkins AT, Das TM, Panzera LC, Bickel SE (2016) Oxidative stress in oocytes during midprophase induces premature loss of cohesion and chromosome segregation errors. Proc Natl Acad Sci U S A 113: E6823–E6830

Perkins AT, Greig MM, Sontakke AA, Peloquin AS, McPeek MA, Bickel SE (2019) Increased levels of superoxide dismutase suppress meiotic segregation errors in aging oocytes. Chromosoma 128: 215–222

Samata M, Alexiadis A, Richard G, Georgiev P, Nuebler J, Kulkarni T, Renschler G, Basilicata MF, Zenk FL, Shvedunova M et al (2020) Intergenerationally Maintained Histone H4 Lysine 16 Acetylation Is Instructive for Future Gene Activation. Cell 182: 127–144 e123

Sinclair DA, Guarente L (2014) Small-molecule allosteric activators of sirtuins. Annu Rev Pharmacol Toxicol 54: 363–380

Singh CK, Chhabra G, Ndiaye MA, Garcia-Peterson LM, Mack NJ, Ahmad N (2018) The Role of Sirtuins in Antioxidant and Redox Signaling. Antioxid Redox Signal 28: 643–661

Spradling AC (1993) Developmental Genetics of Oogenesis. In: The development of Drosophila melanogaster, Bate M., Martinez Arias A. (eds.) pp. 1-70. Cold Spring Harbor Laboratory Press: Cold Spring Harbor, NY

Stunkel W, Campbell RM (2011) Sirtuin 1 (SIRT1): the misunderstood HDAC. J Biomol Screen 16: 1153–1169

Subramanian VV, Bickel SE (2008) Aging predisposes oocytes to meiotic nondisjunction when the cohesin subunit SMC1 is reduced. PLoS Genet 4: e1000263

Tatone C, Di Emidio G, Barbonetti A, Carta G, Luciano AM, Falone S, Amicarelli F (2018) Sirtuins in gamete biology and reproductive physiology: emerging roles and therapeutic potential in female and male infertility. Hum Reprod Update 24: 267–289

Vaquero A, Scher M, Lee D, Erdjument-Bromage H, Tempst P, Reinberg D (2004) Human SirT1 interacts with histone H1 and promotes formation of facultative heterochromatin. Mol Cell 16: 93–105

Vo KCT, Sato Y, Kawamura K (2023) Improvement of oocyte quality through the SIRT signaling pathway. Reprod Med Biol 22: e12510

Wartosch L, Schindler K, Schuh M, Gruhn JR, Hoffmann ER, McCoy RC, Xing J (2021) Origins and mechanisms leading to aneuploidy in human eggs. Prenat Diagn 41: 620–630

Weng KA, Jeffreys CA, Bickel SE (2014) Rejuvenation of Meiotic Cohesion in Oocytes during Prophase I Is Required for Chiasma Maintenance and Accurate Chromosome Segregation. PLoS genetics 10: e1004607

Wu CS, Chen YF, Gartenberg MR (2011) Targeted sister chromatid cohesion by Sir2. PLoS Genet 7: e1002000

Wu QJ, Zhang TN, Chen HH, Yu XF, Lv JL, Liu YY, Liu YS, Zheng G, Zhao JQ, Wei YF et al (2022) The sirtuin family in health and disease. Signal Transduct Target Ther 7: 402

Zeng Y, Li H, Schweppe NM, Hawley RS, Gilliland WD (2010) Statistical analysis of nondisjunction assays in Drosophila. Genetics 186: 505–513

Zhang T, Zhou Y, Li L, Wang HH, Ma XS, Qian WP, Shen W, Schatten H, Sun QY (2016) SIRT1, 2, 3 protect mouse oocytes from postovulatory aging. Aging (Albany NY) 8: 685-696

